# The molecular origin of body temperature in homeothermic species

**DOI:** 10.1101/2024.09.10.612206

**Authors:** Gerhard M. Artmann, Oliver H. Weiergräber, Samar A. Damiati, Ipek Seda Firat, Aysegül Temiz Artmann

## Abstract

We propose the Interfacial Water Quantum-transition model (IWQ model) explaining temperature-dependent functional transitions in proteins. The model postulates that measured critical temperatures, T_C_, correspond to reference temperatures, T_W_, defined by rotational quantum transitions of temporarily free water molecules at the protein-water interface. The model’s applicability is demonstrated through transitions in hemoglobin and thermosensitive TRP channels. We suggest this mechanism also defines basal body temperatures in homeotherms, with T_W_=36.32°C for humans. We demonstrate that human (mammal) and chicken (Aves) body temperatures align with specific reference temperatures, and correlate with pronounced transitions at T_C_ in hemoglobin oxygen saturation. This suggests evolutionary adaptations in homeotherms involve an interplay between oxygen supply and water’s rotational transition temperatures. The IWQ-model states that proteins sense and water sets critical physiological temperatures.

**Significance Statement:** We propose the Interfacial Water Quantum-transition (IWQ) model that offers a new way to understand how proteins respond to critical temperature changes. We suggest that the key temperatures at which proteins change their function are linked to specific quantum transitions of water molecules at the protein-water interface. This model explains why certain critical temperatures, like human body temperature, align with these transitions. By exploring this connection in proteins like hemoglobin and in thermosensitive channels, the IWQ model highlights a fundamental link between water behavior and biological temperature regulation, shedding light on evolutionary adaptations in humans and other animals.

## Introduction

The body temperature of homeothermic species is kept constant within relatively narrow limits by the action of complex regulatory circuits involving thermal sensors and effectors. Despite countless person-years of research, the available body temperature data for homeothermic species, especially humans, remain experimentally determined values that are influenced by many factors (*1–7*). The actual body temperature varies depending on the circadian rhythm, measurement location, state of health, gender and ethnic origin of the subject, etc. (*8–10*). A meta-analysis of body temperatures recorded for 677,423 people, the oldest of whom was born approx. 200 years ago, showed that in the early 19th century the average body temperature of men was 0.59°C higher than it is today, and during this period has decreased monotonically at 0.03°C per decade. For women, the decrease was virtually identical. This trend had been attributed to the improving average state of health of people (*9,10*). It is obvious that this decrease cannot continue in a linear fashion but must approach a threshold value, which may be approximated by the lowest body temperature measured in healthy subjects at rest (36.6°C for males). We refer to this threshold as the reference value of human body temperature. To be considered as such, it must be independent of the above influences; its nature must be determined by a physical mechanism. Such a physical mechanism that could establish reference temperatures for homeothermic species has not been found to date (*11*).

Erythrocytes (red blood cells, RBCs) undergo striking shape changes in the bloodstream due to the prevailing flow forces, occurring within less than 100 ms (*12*). An incidental discovery by Artmann et al. in 1994 initiated the research on the subject of this paper (*13*). If RBCs, with a resting diameter of 7.5–8.7 μm, are aspirated in vitro into a micropipette with an inner diameter of 1.3 µm at a negative pressure of −2.3 kPa (−17.25 mmHg), they block the orifice immediately after entering up to a critical temperature Tc of 36.3 ± 0.3°C (*14*). Above Tc, however, intact RBCs pass through the pipette with almost no resistance (*15*). This transition within ±0.3°C is remarkably sharp and its inflection point at 36.3°C is close to the lower threshold suggested for human body temperature (Fig. 1). This effect was quite a surprise to the RBC rheology and mechanics community (*16–19*).

**Figure 1:**
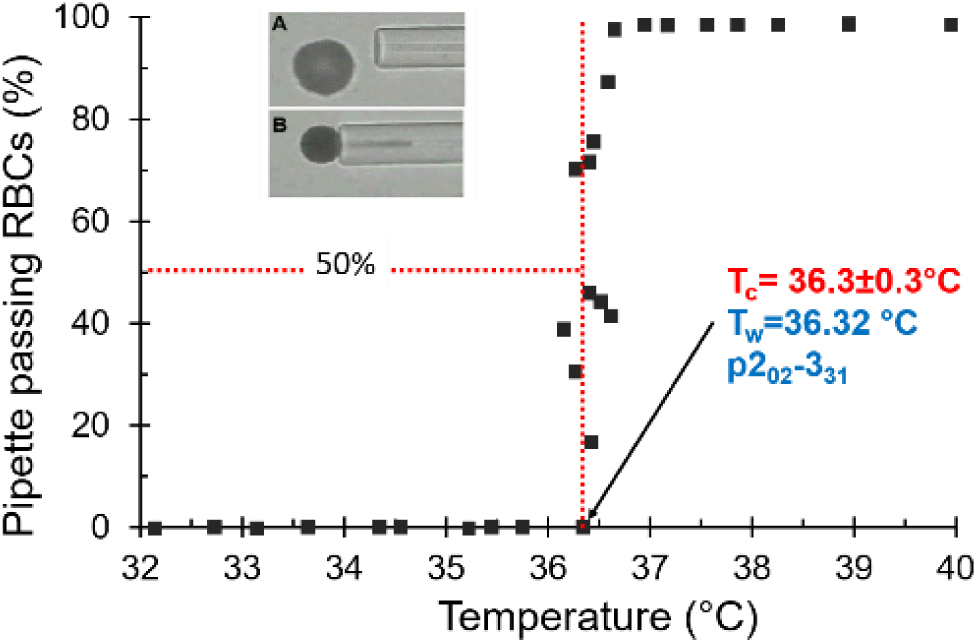
Micropipette RBC passage experiment. Inset photo: Pipette with inner diameter 1.3 µm; RBCs (dark), resting diameter approx. 8 µm, before (A) and during aspiration (B). Graph: Percentage of passing RBCs (squares); Tc, measured critical passage transition temperature; Tw (blue), temperature of the triggering quantum-physical rotational transition of water with spectroscopic nomenclature (see below).

Subsequent experiments conducted in the 30–44°C range, where an initial stage of structural perturbation was found to occur in hemoglobin, aimed to understand the cause of the jump-like temperature transition of RBC passage (*20,21*). Our interdisciplinary effort involved a wide variety of methods, such as the micropipette aspiration technique low shear rotation viscometry, circular dichroism (CD) and dynamic light scattering measurements as well as NMR spectroscopy and various neutron scattering experiments on intact RBCs and hemoglobin samples (*21–23,26–28*). In particular, CD measurements revealed that hemoglobin of all homeothermic species studied undergo a partial and reversible thermal transition in the structural perturbation range with turning points, Tc, close to the respective species’ body temperatures (*25*). We concluded that hemoglobin acts as a molecular temperature sensor (*23*). Temperature transitions with Tc between 36°C and 37°C were found with all experiments with human RBCs or hemoglobin, indicating that Tc was neither a cellular nor an RBC membrane effect; moreover, it was independent of the pH of the buffer (range 6.8–7.8) and the calcium concentration (0–18 mM) (*22,29*). Intriguingly, temperature transitions were observed not only for oxy- and deoxy-hemoglobin but also for methemoglobin and even sickle cell hemoglobin and thus appear to be inscribed into the globin architecture (*21*). In NMR T_1_ measurements of highly concentrated RBCs in autologous blood plasma versus blood plasma alone, we found a transition at Tc approx. 37°C in both samples, but the temperature dependence of T_1_ surprisingly showed opposite trends on either side of Tc (see Fig. 3 in *23*). Finally, micropipette passage experiments with human RBCs in D2O-based buffer showed a passage jump at Tc=37.2°C, i.e. 0.7°C higher than in H_2_O-based buffer (see Fig. 5 in ref. *23*), pointing to a role of the solvent “water” in this transition (*30,31*).

The body temperatures of homeothermic species are subject to evolution (*8, 32–35*). Indeed, the temperature transitions of hemoglobin are species-specific (see Fig. 4 in ref. *25*) (*24,36*). The RBC findings per se are mainly of interest to specialists; however, in-depth research into the physical nature of this phenomenon has led us to the fundamental concept that water is the key to understanding the observations. As a mechanism, we postulate that at the critical transition temperatures a molecular switch is triggered, which is based on the interaction of hemoglobin with water in the protein-water interface. This postulate does not require any specific assumptions about the structure of hemoglobin and should therefore be of general significance, i.e., be applicable to other proteins (*22,31,37–39*).

In homeothermic species, the hypothalamus is in charge of actively regulating body temperature (*40*). It obtains information on local temperatures in the organism via thermosensitive neurons with afferent cold and warm fibers (*5,41,42*). The actual temperature sensing, i.e., the conversion of the physical variable temperature into electrical signals, is carried out by membrane-spanning, temperature-sensitive cation channels that belong to the **t**ransient **r**eceptor **p**otential (TRP) channel superfamily and feature exceptionally high temperature coefficients (Q_10_). Due to molecular adaptations, certain TRP channels are specifically „heat-activated” while others are „cold-activated“. While the precise molecular mechanisms governing temperature sensitivity are still largely unknown (*43–49*), there are numerous indications that the protein-water interaction in the water exposed parts of TRP channels may play a key role (*45,50–58*). As is the case with hemoglobin, the temperature dependence of TRP channels is expected to differ from species to species (*11*).

In this work, we propose the interfacial water quantum transition (IWQ) model as a novel paradigm for the explanation of temperature-dependent structural and functional transitions that occur at critical temperatures in proteins and other biomolecules. Its central postulate is that the experimentally measured transition temperatures, T_C_, are determined by quantum-physical reference temperatures, T_W_, relating to rotational transitions of temporarily free water molecules in the protein-water interface (*59–65*). After introducing the concept and explaining the transitions observed in RBCs and hemoglobin, we turn to membrane proteins, mainly thermosensitive TRP channels. Based on the rapidly growing body of knowledge on their structure and function, we use the IWQ model to justify the characteristic temperatures at which their thermal sensitivities switch between different Q_10_ regimes (*43,45,47,49,55,56,58,66*). Finally, we explain why the body temperatures, T_B_, of homeothermic species are always restored in the process of thermoregulation and do not drift on the temperature scale. Using previously unpublished data for human (mammalian) and chicken (avian) hemoglobin, we show that the above-mentioned transition at the body temperatures of the respective species expresses itself as an inflection point in oxygen saturation. This suggests that the evolution of body temperatures in homeotherms may have been dominated by the optimisation of critical physiological parameters in light of the interplay between available quantum-physical rotational transitions of water at T_W_ on the one hand and the peculiarities of the protein interfacial structure on the other.

## The Interfacial Water Quantum-Transition Model (IWQ Model)

### A thermally expanding, spherical model protein in aqueous solution

Water is the most important solvent in biological systems and is fundamental to life (*64,67–71*). To set the stage for introduction of the IWQ model, we need to consider several important aspects of the anisotropic thermal expansion of a globular model protein (e.g. hemoglobin, myoglobin, serum albumin, immunoglobulins) in aqueous solution (*72–74*). In an aqueous, physiological environment, the majority of apolar amino acid residues are sequestered in the protein core due to their hydrophobicity and are not very dynamic. The mainly hydrophilic, outer side chains form a protein-specific surface landscape and, together with the adjacent water, constitute the overall protein-water interface. The global thermal expansion coefficient of myoglobin crystals is 115·10^−6^ K^−1^ (between 255 K and 300 K), for liquid water it is 70·10^−6^ K^−1^ (to compare and contrast, water ice 5·10^−6^ K^−1^, glass 8·10^−6^ K^−1^). Hence, myoglobin thermally expands about twice as much as bulk water (*75*). The thermal expansivity of proteins is subject to large variability and is highly anisotropic, which is essentially due to the influence of non-hydrated internal cavities, i.e., a variation in the density of internal vdW interactions.

### Topography and dynamics of the protein-water interface

The distribution of hydrophilic and hydrophobic surface areas of the protein is temperature-dependent due to the anisotropic thermal expansion of near-surface protein structures. As the thermal energy increases, secondary and tertiary protein structure elements tend to be destabilised, and distinct structural transitions can occur. The process is typically sigmoidal over temperature. At the same time, the dynamics of the protein-water interface increases (*20,21,76–79*). In *polar* surface areas, non-uniformly distributed hydrogen bonds form an irregular network between the protein and adjacent water. They bind and break continuously, only some are long-lived. In contrast, *non-polar*, hydrophobic surface areas are covered by clusters of water molecules that are not hydrogen-bonded to the protein (Fig. 2A), which have been termed cavity-wrap water (*79,80*). The resulting topological profile is dynamic with time constants according to Pal et al. of 16-43 ps (*71*).

**Figure 2A:**
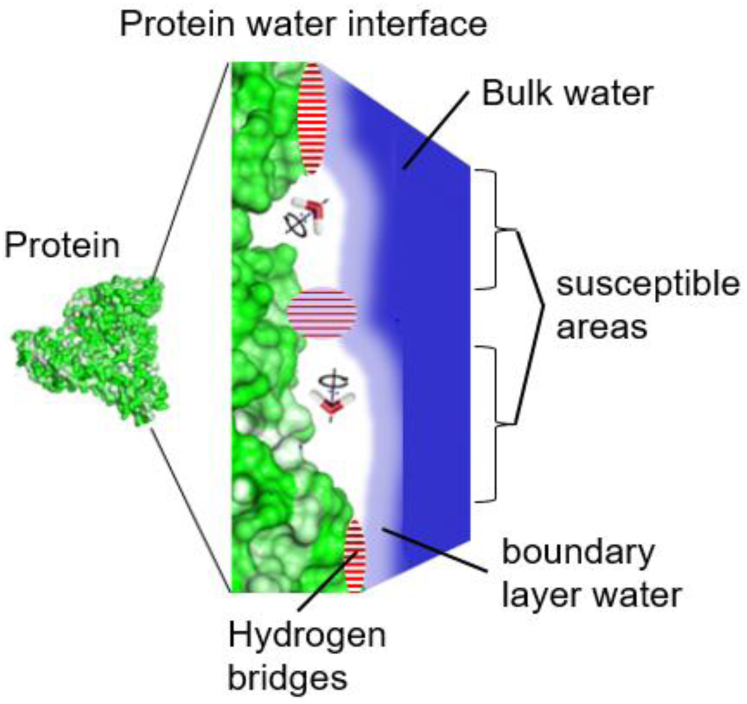
Topography of a protein-water interface. left, surface representation of a generic protein; right, close-up illustrating protein-solvent interactions. Hydrophilic surface regions are involved in protein-water H-bonds with an irregular internal density (dashed red). In the hydration shell covering the surface area (light to dark blue), translational diffusion of water is restricted, and both density and order of hydration water is greater than that of bulk water. Hydrophobic surface areas (“susceptible” areas, see below), which may enlarge or newly arise during thermal expansion, are proposed as hot spots of rotational water transitions (81,82).

During thermal expansion, additional hydrophobic components are exposed on the protein surface, with ensuing changes in geometry and hydration (*37,83,84*). The transfer of bulk water to newly-exposed (hydrophobic) surface areas requires inter alia an osmotic work of solvation, which contributes to the energetics of protein and solvent restructuring (*26,28,37,70,85*).

The translational and rotational dynamics of water molecules is generally slowed down in the interfacial region due to hydrogen bonding with protein residues as well as volume exclusion effects. It seems plausible to assume that, particularly at hydrophobic sites devoid of hydrogen bonds between protein and water, the dynamics mismatch may result in temporary emergence of regions with reduced density of solvent molecules. This intuitive view is supported by simulation studies investigating the dynamics of water in hydration layers. Unlike their hydrophilic counterparts, hydrophobic molecular surfaces are characterized by deviations from the Gaussian distribution of hydration water densities, featuring low-density tails. Indeed, the free energy required to form a solvent-free cavity (ΔG_cav_) has been proposed as a generally applicable proxy to quantify hydrophobicity, even for surfaces as complex as those of proteins (references therein *37*).

### Rotational quantum transitions of free water molecules

Water exists in two isomeric forms, as para-water (p-water) with the proton spin J=0 and ortho-water (o-water) with J=1 (Fig. 2B). In their unbound (gas-like) forms, both isomers undergo quantum-physical, molecular rotations involving distinct states that are described by quantum numbers (*62,65,66,86–90,*).

**Figure 2B:**
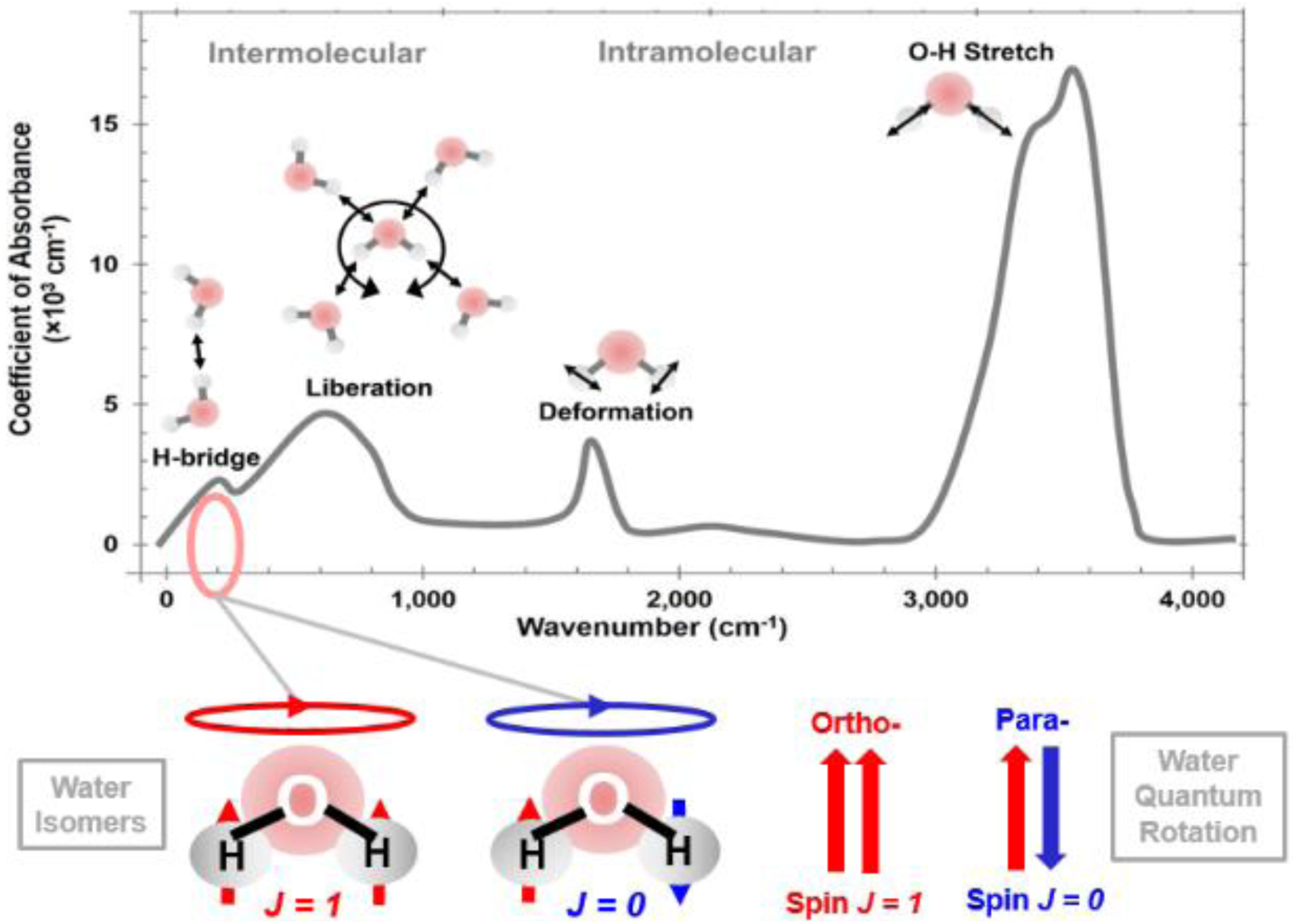
**Above**: Scheme of infrared absorption spectrum showing different vibrational modes of water molecules in the THz-infrared region. Marked (pink oval) is the energy range of the rotational transitions considered herein. **Below**: Para p-water (non-magnetic) with spin J=0 and ortho o-water (magnetic) with spin J=1.

Transitions between rotational states occur by absorbing or releasing precisely defined quantum energies in the terahertz range of the electromagnetic spectrum (Fig. 2B, C). Due to the role of spin conservation, spontaneous transitions of p-water into o-water isomers and vice versa are quantum mechanically forbidden. An exception is when the energies of an o- and a p-level are close together (green dashed line, ΔE= 0.2 cm^−1^).

**Figure 2C:**
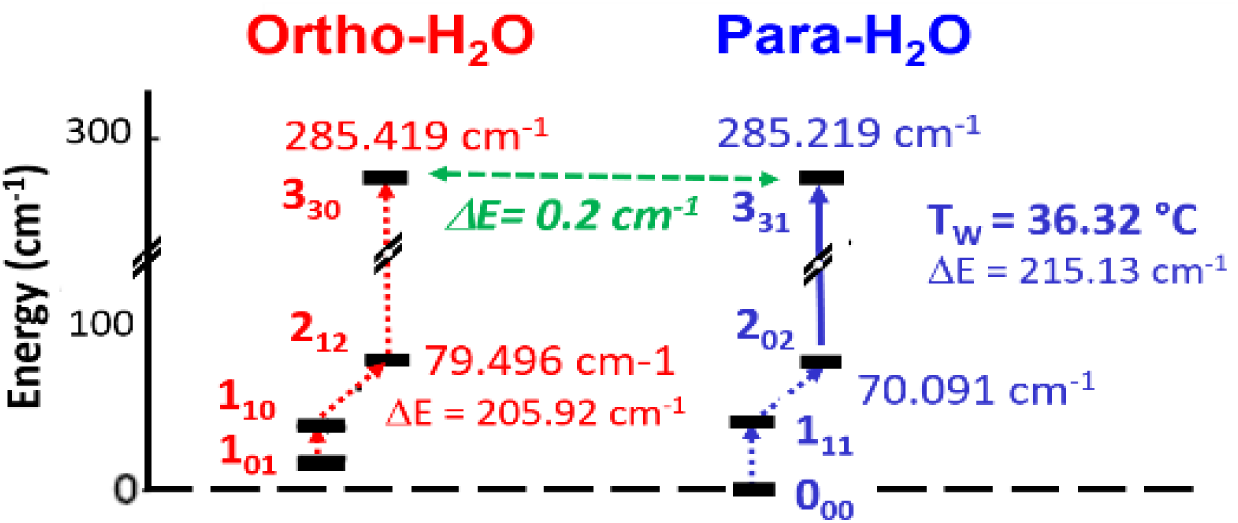
Excerpt from the rotational quantum transition scheme of water isomers (86,87,91). The ground state energy of p-water (blue), p000, is zero; it does not rotate in this state, while the ground state energy of o-water, o101, is greater than zero and always rotates (red).

Spin conversion is as well facilitated by the close proximity of some biologically important isotopes acting as catalysts. Its rate depends on catalyst’s magnetic field gradient and density. Catalysts are for example triplet ^3^O_2_, ^57^Fe (2.1%), ^39^K (44%), ^23^Na (100%), ^67^Zn (4%), ^25^Mg (10%), ^43^Ca (0.135%), ^13^C (1.1%), ^31^P (100%) (*92,93*).

In the past, the analysis of rotational spectra was largely limited to gases in which water molecules can rotate freely (Figure 2A) (*60*). While an analogous situation can be generated by embedding water isomers in C_60_ fullerene cages, in which they cannot form hydrogen bonds (*94–97*), rotational transitions are commonly thought to be suppressed in bulk water because of the extensive hydrogen bonding network (*62,98*). This fundamentally correct view has been relativized by experiments using DNA and protein solutions (*65,91,99*) (discussed in the following chapter) as well as recent quantum mechanical MD simulations. The latter highlight the statistical distribution in the density of hydrogen bonds in liquid water: In addition to the highly populated states (three or four H-bonds per water molecule), there are also states with lower coordination numbers. In fact, around 2% of all water molecules in pure water were found *unbound* at any given time, which would allow for short-lived quantum rotation without calling into question the fundamental statement on the extensive suppression of such transitions (*100,101*). Moreover, the hydrogen-bonding network is well known to get more dynamic as temperature increases, with an anticipated growth of the fraction of instantaneously free water molecules. Adding to the fluctuations of hydrogen bonding density in bulk water, hydrophobic patches on solvated macromolecules are likely to constitute hot spots for water quantum rotation, due to the enhanced probability of low-density states favoring the occurrence of free water molecules (*99*). Effective hydrophobicity is assumed to be particularly prominent in high-curvature concave surface elements, suggesting that deep and narrow apolar cavities should provide the highest probability of vapor-like water (*82*). These considerations support the view that both in pure water and in the protein-water interface (Fig. 2A) a small fraction of shortly “free” water molecules is available for rotational quantum transitions.

### Experimental evidence of rotational transitions in aqueous biopolymer solutions

Rotational transitions in gaseous water can be detected with Raman spectroscopy and other classical methods of infrared spectroscopy (*78*). However, they are not well suited for resolving rotational transitions in liquid water or the protein-water interface with sufficient sensitivity because of the low proportion of usable signal, the unfavorable signal-to-noise ratio and the considerable absorption of the water (*61,63,99,102,103*).

For the detection of rotational transitions in liquid water, significant progress has been made with coherent laser spectroscopy methods. These methods are characterized by high spectral resolution and an excellent signal-to-noise ratio (*61,102,104–107*). This is due to the fact that the rotational spectra of the two water isomers are more dependent on the molecular mass than vibrational spectra, which are normally used to study molecular interactions in liquids (*65,103*). Using coherent laser spectroscopy, rotational transitions in pure liquid water as well as in aqueous solutions of DNA and protein have been demonstrated experimentally (*99,103,108*). It is important to note that the spectra of biomolecular solutions differ from those of pure water, confirming the notion that the protein or DNA surface provides sites that promote quantum transitions of the solvent. The peak amplitudes in their rotational spectra are a measure of the number of rotational transitions of free water molecules and their interaction with (susceptibility to) the electric field of two coherent laser beams at a given wavenumber. Peaks (resonances) appear in the spectrum at wave numbers λ^−1^ inducing rotational transitions in the sample volume (*102,106*). In these experiments, the rotational spectrum of water isomers has thus emerged as a spectroscopic fingerprint of the dissolved biomolecule.

### Mechanistic foundations of the IWQ model

The local and temporary occurrence of free water molecules without H-bonds in the protein-water interface is a fundamental premise of the IWQ-model. In addition, given that the temperature of water (or any kind of matter) is directly related to the internal energy of its constituents, we adopt the view that rotational transitions can be initiated as characteristic resonance temperatures, TW, are reached (*109,110*). Within the temperature range of protein structural perturbation, experiments often reveal transitions at critical temperatures, TC, at which protein functions or other properties change discontinuously (see below for examples). According to the IWQ model, these transitions can be explained by energy uptake of free water molecules in the protein-water interface (Fig. 2A). These may undergo rotational quantum transitions by absorbing the discrete energy ΔE_n,m_ that is defined by the rotational quantum states n and m, respectively. Assuming that such processes are initiated by collisions with thermal energy k·T_W_ ties each transition to a characteristic rotational transition temperature TW

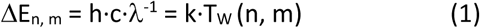

with h = Planck’s constant, c = the speed of light, k = Boltzmann’s constant, and λ^−1^ = the wavenumber (*62,109*). All physically possible rotational transitions of free water involving quantum states of lower energy difference have already taken place before a specific TW is reached, thus contributing to the energy landscape of the protein surface below T_W_ (*81*). At T_W_ and above, however, absorption of higher quantum energies becomes possible, with effects determined by the actual number and distribution of free water molecules in the interface (Fig. 2A). When the excited water molecules release their energy and re-integrate into the hydrogen-bonding network, this leads to local energy deposition at the protein-water interface. We propose the term “susceptible areas” for regions of the protein-water interface where rotational transitions are possible. At any given temperature, there may be several such areas, the size and distribution of which are protein-specific (*111,112*). If the water-mediated energy input at T_W_ into these susceptible areas is sufficiently high, the protein can undergo local structural changes, thus attaining a new conformational equilibrium, which is often associated with a switch in protein function.

If the energy, often determined as enthalpy change ΔH, taken up by the protein during a temperature-dependent transition at Tc is known, the number of water molecules required to deliver this amount of energy via rotational transitions may be estimated from the quantum energies involved (*86,87*):

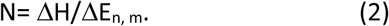

As the temperature continues to increase, further rotational transitions are excited. Due to the changes in surface properties of the protein, at each rotational transition temperature a partially or completely different set of susceptible regions can be addressed, resulting in further conformational changes. The concept of susceptible surface areas not only applies to soluble proteins but also to membrane-spanning ones. Obviously, in these cases local energy accumulation via water quantum transitions is restricted to solvent-exposed regions or domains (Fig. 2A, and 4E, F).

The IWQ-Model postulates that, in the range of reversible perturbation, many pronounced temperature-dependent transitions in protein structure and function cannot be explained by continuous energy uptake of the protein-water system with rising temperature but require local energy input after a discrete water rotational transition temperature T_W_ is reached. Thus, T_W_ may represent the physically defined reference temperature of the protein transition observed at TC. Since T_W_ is based on a quantum-physical property of water molecules, it is independent of solution pH, chemical potential, salt concentration and other proteins in the aqueous environment.

What happens as the temperature decreases? When the temperature drops below T_W_, the energy introduced into the local protein interface regions is recovered. In the structural perturbation stage, IWQ-transitions should be reversible, unless this is hindered by secondary reactions, such as protein aggregation or semi-permanent thermal denaturation, as with some temperature-sensitive TRP channels (*21,23,79,113,114*).

## Application of the IWQ model to human RBCs and hemoglobin

### Temperature transition of RBC passage through micropipettes (RBC passage transition)

The cytosol of RBCs is mainly comprised of freely suspended, almost spherical Hb molecules that form a crowded medium (*27,115–117*). It exhibits a Newtonian flow behavior at physiological concentration (*12*). The relaxation times of RBCs, which are determined not only by the cell membrane viscosity but also by the viscosity of the cytosol, decrease linearly and monotonically with increasing temperature up to ∼ 40°C (*19,118*). In particular, it does not show anomalies around human body temperature (*119–121*).

In contrast, passage experiments with RBCs suspended in phosphate-buffered saline through 1.3-µm micropipettes at a high aspiration pressure of −2.3 kPa revealed a very sharp and highly reproducible temperature transition in the passage ability (*RBC passage transition*) at T_C_= 36.3 ± 0.3°C (*Fig. 1*). Below T_C_, the RBCs block the pipette, while the Hb concentration in the spherical RBC trail remaining outside the pipette increases to 50 g/dl (Fig. 3A) (*13*). From ∼37°C onward, however, they pass through unhindered (*15*).

**Figure 3A:**
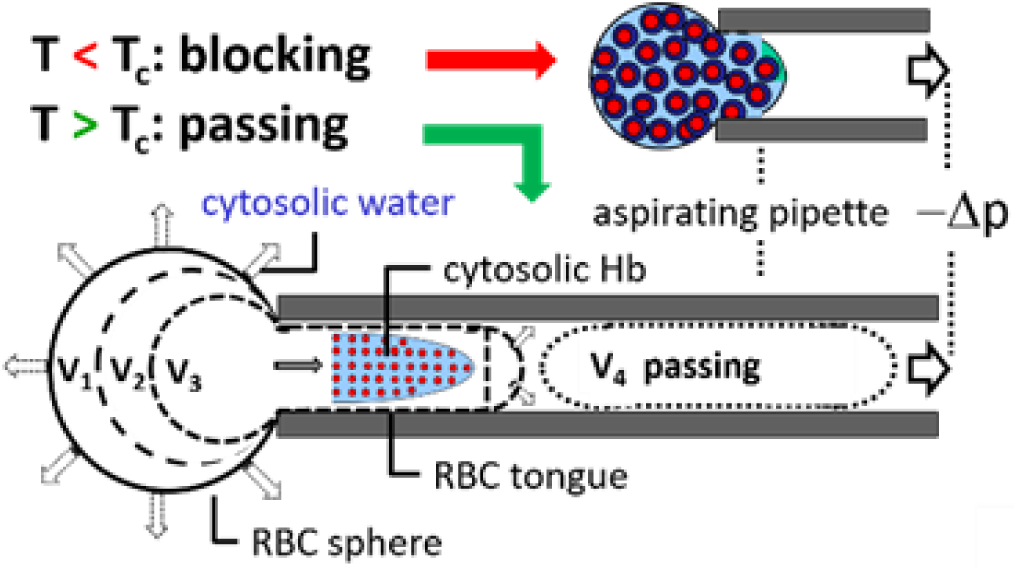
RBC passage transition. Human RBCs in a micropipette undergo a step-like transition at the critical temperature T_C_= 36.3 ± 0.3°C. During the entry process, the Hb solution in the RBC cytosol is mechanically sheared. Simultaneously, water is displaced from the cytosol, reducing the volume of the outer RBC sphere. Inside the sphere, Hb forms a highly concentrated, viscous Hb-gel. At T_C_, its viscosity undergoes a phase transition, allowing the whole RBC to pass the pipette. At T>37°C, the pipette does not cause any significant resistance to the RBC passage.

Applying the IWQ model, we postulate that this is the result of quantum mechanical rotational transitions in the Hb-water interface at T_W_=36.32°C (p2_02_-3_31_). This transition causes a sudden drop in cytosolic fluidity. Within the accuracy of the measurement, the temperatures T_C_ and T_W_ can be considered identical; moreover, they correspond to the average body temperature of a healthy person, T_B_ = 36.6 °C (99% range 35.3-37.7°C) and thus might actually represent the basal limit of the human body temperature (*9,10*).

To test whether the isotopic state of the hydrogen in the water molecule has an influence on the RBC passage transition, micropipette experiments were performed with RBCs suspended in phosphate buffer containing Deuterium dioxide, D_2_O, instead of H_2_O. Deuterium-based hydrogen bonds are slightly stronger than protium-based ones due to the difference in mass between deuterium (D) and protium (H), implying a higher stability of protein-solvent and solvent-solvent interactions. Thus, it was expected that the critical transition temperature should be shifted to higher values, which we indeed observed. The critical transition temperature for RBC passage was T_C_ (D_2_O) =37.2°C, corresponding to an upward shift by 0.8 °C. In accordance with the IWQ-Model, we assign this inflection point to the rotational transition p9_82_-9_91_ with T_W_=37,09°C (see Fig. 5 in *23*).

### Temperature transition of RBCs trapped in micropipette orifices (RBC volume transition)

With the same negative pressure in the pipette as above, but a smaller internal diameter of 1.1 µm, at which no RBC can pass through, the temperature dependence of the equilibrium RBC volume, ΔV/ΔT, between 34.5°C and 39.5°C was measured (Fig. 3B) (*23*). Since the RBC membrane remains intact during pipetting, the aspiration force causes an increase in cytosolic pressure.

**Figure 3B:**
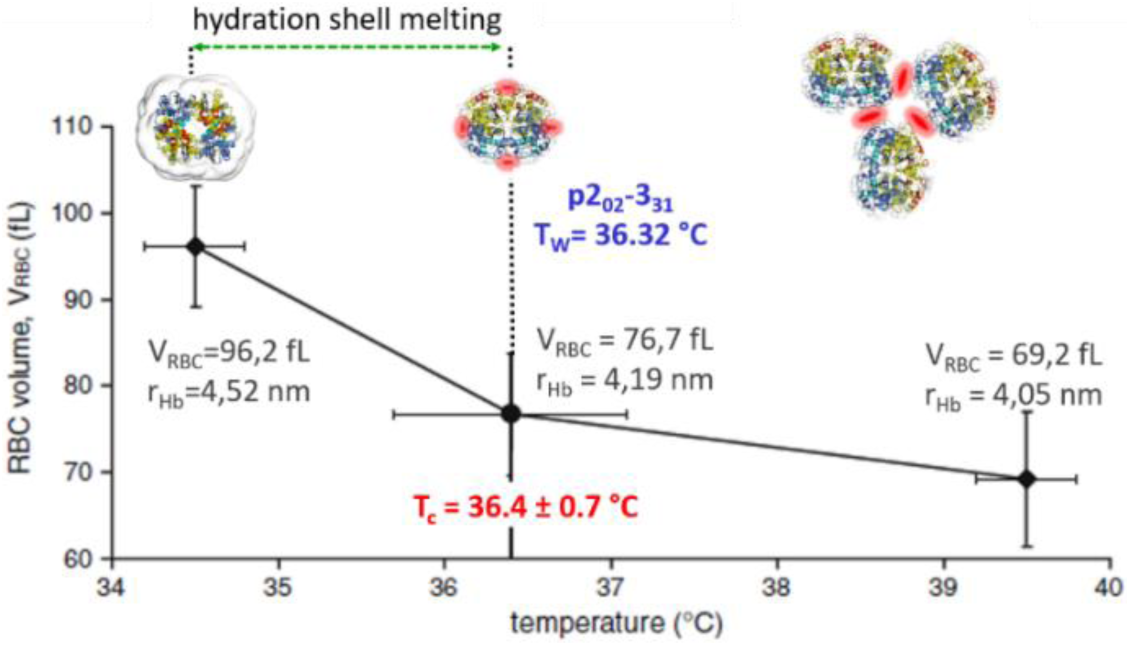
RBC volume transition. (with kind permission from Artmann et al. 1998 (13). Individual RBC volume as a function of temperature is shown. For each temperature step 12 individual RBCs were aspirated. Single RBC volume was calculated from microscopic images. A t T_C_=36.4±0.7°C there was a significant kink. Below T_C_, the volume loss slope was 4.2 times higher than above. Apparently, water “melted” from the Hb hydration shell at a given temperature into bulk water is expelled from the cytosol. Above the TC, the rate of water molecule detachment from the hemoglobin surface is minimal.

Consequently, the cytosolic bulk water is squeezed out through the aquaporin channels*67* until an equilibrium is reached, which is presumably dictated by the amount of water tightly bound by hemoglobin, the major colloid osmotic agent in the RBC cytosol. The observed decrease in RBC volume with increasing temperature can thus be attributed to the conversion of Hb-bound water to bulk water. In other words, at this temperature interval, the HB-hydration shell “melts” and bound water becomes bulk water (*23*). At T_C_=36.4±0.7°C an inflection point of the equilibrium RBC volume was found (*RBC volume transition*), again coinciding with the rotational transition p2_02_-3_31_ at T_W_=36.32°C. The volume loss with temperature (ΔV/ΔT) was quite steep at T<T_C_ (−10.3 fl/°C) but flattened to −2.4 fL/°C above Tc, indicating that the melting process of Hb-bound water is essentially complete at T_C_=T_W_.

As further indirect evidence for the loss of water bound to Hb below T_C_, we note that in NMR measurements on plasma-suspended RBCs the T_1_ relaxation time increases in this temperature range but remains almost constant above the transition temperature (see Fig. 5 in *23*). It is crucial to understand that the *RBC volume transition* is physically distinct from the *RBC passage transition*. While the former is a stationary phenomenon, the latter is a fluid-dynamic effect in which shear forces act.

Estimates showed that the applied thermal energy is too small to explain the inflection points of the two pipette experiments (Fig. 3A and B). The small difference in colloidal osmotic pressure of the cytosol in the selected temperature range cannot have caused the transition either (*22*). Thus, in our opinion, classical physicochemical theories are unable to explain the occurrence of the observed inflection points. Together with temperature-dependent dynamic light scattering experiments and Hb-solution viscosity recordings (Fig. 3C) the above data support the view that the rotational transition of water at T_C_ leads to a sudden increase in hydrophobicity of the Hb surface, enabling low-friction sliding of the Hb molecules and hence the RBC passage transition with the 1.3-µm pipette (*13,24,25*). Note that we are referring to an *effective* hydrophobicity, which cannot simply be deduced from the fraction of apolar moieties exposed on the protein surface. In fact, the numerous solvent-accessible side chain and main chain atoms featuring different polarity cooperate in complex and non-trivial ways, depending on their relative arrangement, to yield local and global patterns of hydration (*37,82,122*). Consequently, a moderate rearrangement of a given set of protein surface elements may be sufficient to alter solvent interaction, and even though an increase in temperature may be expected to result in partial unfolding and hence exposure of additional apolar side chains, the effect on global hydrophobicity may be disproportionate. The mechanism we are proposing for the *RBC passage transition* differs fundamentally from a previously expressed view (*110*).

**Figure 3C:**
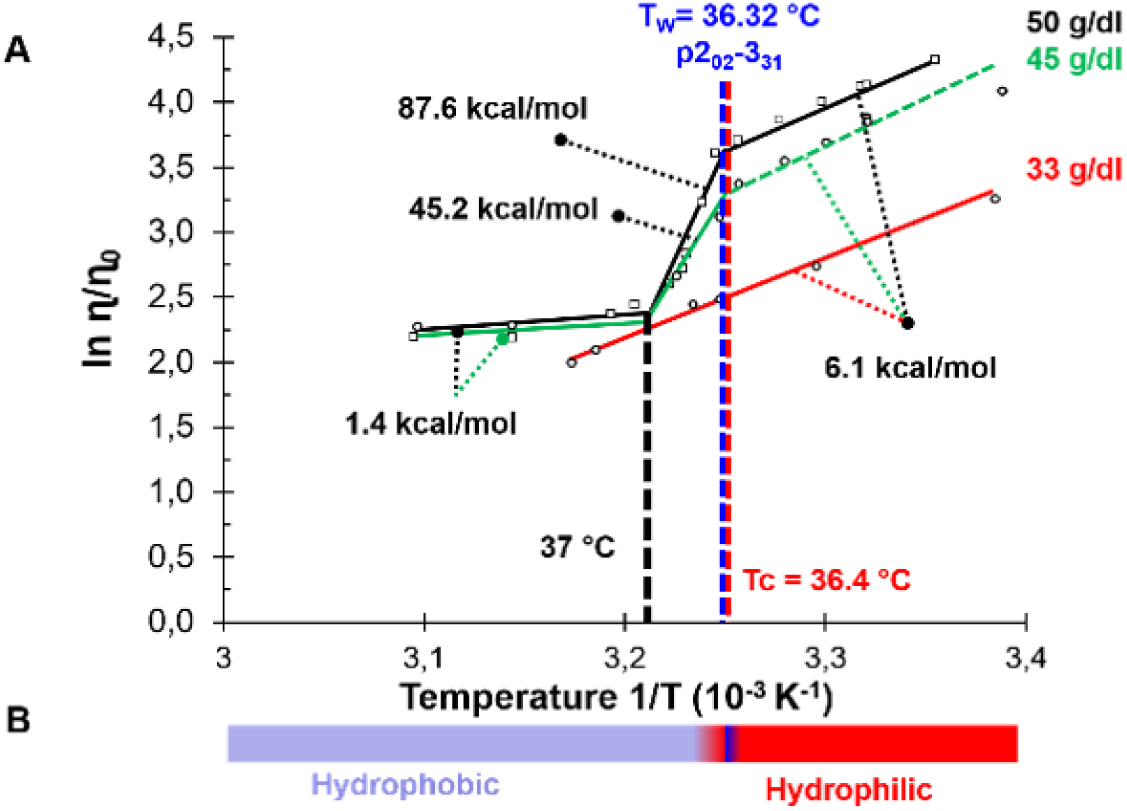
Hemoglobin viscosity transition. (with kind permission from Artmann et al. 1998 (13): Arrhenius plot of the relative viscosity of concentrated human hemoglobin. ***(A)*** At 33 g/dl, the viscosity decreases monotonically with increasing temperature. No temperature transition was seen. At 45 and 50 g/dl, however, a transition became visible at TC=36.4±0.3°C corresponding to the IWQ transition p2_02_-3_31_ at T_W_=36.32°C. The relative viscosity is concentration-dependent at temperatures below T_C_. From T ∼ 37°C on, there was no difference between the two higher concentrations within measurement accuracy.***(B)*** The viscosity transition of hemoglobin (Hb) is attributed to a sudden change in effective hydrophobicity at the Hb surface.

The narrower 1.1-µm pipettes (Fig. 3B) do not allow for any RBC passage but uncover a release of bound water to the bulk of the cytosol, which precedes the actual transition at T_C_. The intriguing observation that melting of the hydration layer ends (within the precision of the experiment) at the transition temperature, T_C_, suggests that the loss of hydration water may sensitize the protein to the energy uptake by quantum absorption, e.g. by altering the dynamics of surface-exposed residues, which has been observed as well in neutron scattering experiments (see Figure in 26).

## Temperature transitions of Hb solutions in a rotary viscometer (Hb viscosity transition)

To rule out RBC-membrane-related switches, such as a sudden loss of adhesion between the RBC membrane and the inner surface of the pipette or a sudden change in the water permeability of the aquaporins, as causes of the RBC transitions described, concentrated Hb solutions were analyzed with a rotational viscometer at a shear rate of 5.89 s^−1^. The equilibrium time between successive temperature steps was 15 min. Hb concentrations were chosen that occur in the RBC sphere (Fig. 3A) during the micropipette *RBC passage* (Fig. 3C) (*13,22,29*). Considering the IWQ-Model, the observed transition at T_C_=36.4°C is caused by rotational transitions p2_02_-3_31_ of free water molecules at the Hb-water interface. At approx. 37 °C, a stall occurs, which leads to the observed independence of the viscosity of the Hb solution from the concentration at elevated Hb. The viscosity range between 36.4 and 37°C represents a transition state.

It is important for the comparison and interpretation of the data that fluid dynamic shear forces act in the cytosol or in the Hb solution in both Micropipetting experiments (Fig. 3A) and rotational viscometry (Fig. 3C). In contrast to viscometry, however, in the pipette experiments, cellular water is expelled into the extracellular space at the same time. The total volume of the RBC is reduced, which implies that the average water content per molecule of Hb is also reduced. The Hb concentration in the RBCs globular part remaining outside the pipette increases up to 50 g/dl below T_C_= 36.4 ± 0.3°C (*13,23,123*). Our observation of hydration water melting off Hb in RBCs as a function of temperature must also hold in the viscometer, but unlike in RBCs under aspiration stress, the average water content per Hb molecule remains constant (*22*).

The relative viscosity, ln (η/η_0_), in the Arrhenius plot decreased monotonically and linearly with 1/T between 23°C and 42°C for the 33 g/dL Hb solution. There was no inflection point. In contrast, at Hb concentrations of 45 and 50 g/dl, an abrupt and significant temperature transition in Hb relative viscosity was observed at T_C_=36.4°C (*Hb viscosity transition*) (Fig. 3C). Up to T_C_, Hb solution viscosity was concentration dependent, in agreement with expectation and with published data^120^. The ratio of free bulk water molecules, N_free_, to hemoglobin-bound water molecules, N_bound_, at T approx. 25°C and concentrations of 33 g/dl, 45 g/dl, and 50 g/dl is estimated to be N_free_/N_bound_= 5.3, 3.0, and 2.0, respectively, corresponding to approx. 7, 4, and 3 molecule layers of *free bulk water* between adjacent Hb molecules (*22*). Despite these differences, the Arrhenius plot yields the same apparent activation energy, within the acuracy of the measurement, in all three cases (6.1 kcal/mol). Intriguingly, the Hb viscosity transition is only observed when the translational diffusion is severely restricted due to the close proximity of the Hb molecules. The jump in activation energy at T_C_=36.4°C amounts to ΔE_a_= 45.2-6.1 kcal/mol = 39.4 kcal/mol at 45 g/dl, and to ΔE_a_ = 87.6-6.1 kcal/mol = 81.5 kcal/mol at 50 g/dl. At approx. 37 °C the temperature dependence of relative viscosity changes again for the higher Hb concentrations, switching to an apparent activation energy of only 1.4 kcal/mol; this corresponds to a difference ΔE_a_ = 1.4-6.1 kcal/mol = −4.7 kcal/mol w.r.t the low-temperature regime.

Applying the IWQ-Model, we propose the increase in Hb hydrophobicity resulting from water quantum absorption at T_W_=36.32°C to trigger a fluid dynamic instability in concentrated Hb-solutions (*124*). The first hydration shell in the Hb interface contains approx. 2300 water molecules reported by Stadler et al., 2008 (*27*). Since the energy absorption of the rotational transition of a water molecule at p2_02_-3_31_ is known, the percentage of water molecules that are required to provide the energy for the viscosity jump can be estimated from the total energy taken up by the Hb molecule including its hydration shell. Taking the activation energy differences mentioned above as a proxy yields numbers of 64 and 133 (Equation (1) and (2)) for the transitions at 45 and 50 g/dl, respectively, amounting to only 2.8% and 5.8% of the water molecules of the first hydration shell.

The ultimate cause of these temperature transitions is water. Even after calcium-induced crosslinking of cytosolic hemoglobin in RBCs or hemoglobin in solution, we found a temperature transition at a critical temperature T_C_ = 36.4 ± 0.3°C which again coincides with T_W_ = 36.32°C of the rotational transition p2_02_-3_31_ (*22,29*). This means that the position of T_C_ on the temperature scale remained unchanged even after significant alteration of protein-protein interactions. As expected, calcium did have strong effects on RBC passage through micropipettes and Hb suspension viscosity, confirming that environmental factors influence the biological effects induced by rotational transitions of water. The transitions themselves, however, always take place at the same temperatures, which are predetermined by quantum physics.

### Hemoglobin Solution Circular Dichroism (Hb CD transition)

In the temperature range between 35°C and 39°C, CD measurements with human hemoglobin showed a transition in ellipticity at 222 nm, indicating an accelerated decrease in α-helix content (*125*). Independent measurements on different CD instruments resulted in transition temperatures of Tc = 37.2 ± 0.6°C, T_C_ = 37.1 ± 0.5°C and T_C_ = 36.5 ± 0.5°C (*21,24,25*). We attribute the varying T_C_ to systematic deviations of the actual temperature in the sample volume from the temperature set on the device. As reported by Artmann et al., the CD transition is independent of Hb oxygenation, pH and other factors (Fig. 3D) (*21*).

**Figure 3D:**
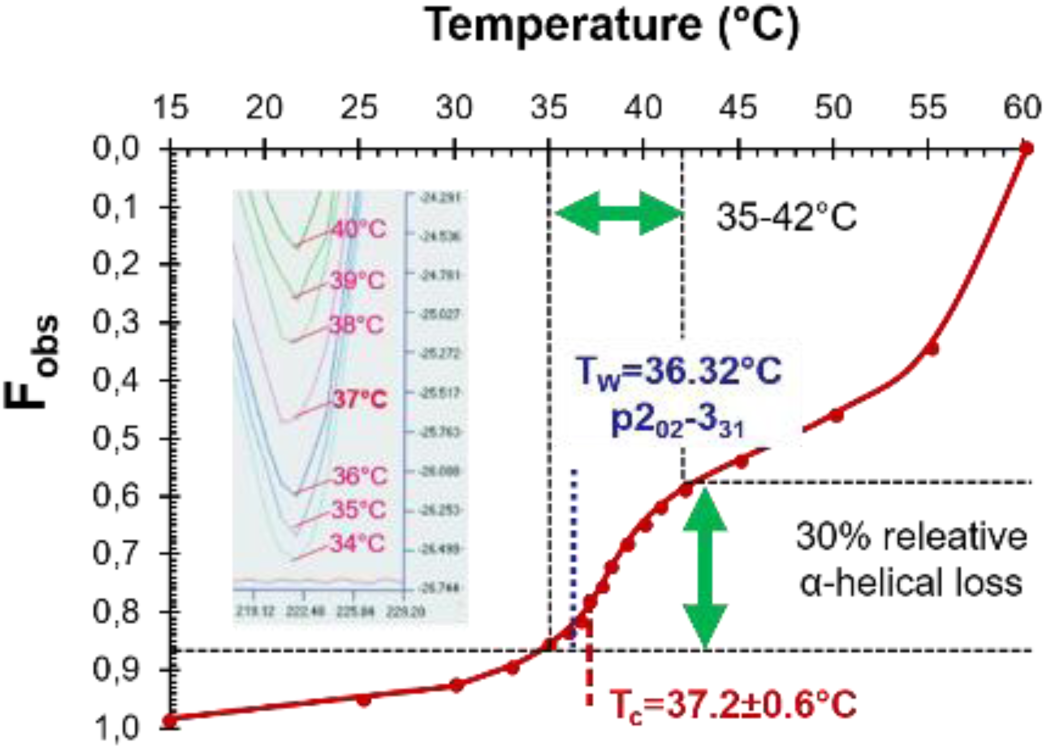
Hemoglobin circular dichroism transition. (reproduced with kind permission (21): Fractional ellipticity change at 222 nm (F_obs_) (21) with temperature for human hemoglobin at pH steps between 6.8-7.8 (Inset: Original CD-Plot, pH 7,4).

We assign the rotational transition p2_02_-3_31_ of water at T_W_=36.32°C to these critical temperatures. In accordance with the IWQ model and our conclusions for other Hb-related transitions, this accelerated decrease in α-helix content correlates with an increase in Hb surface hydrophobicity. This can be inferred from independent dynamic light scattering measurements on human Hb revealing T_C_= 36.4±0.8°C and T_C_= 36.5±0.5°C (*24,25*). Above these T_C_s, the aggregation of Hb increases at a significantly elevated slope. The CD transition is therefore likely to be based on the same quantum mechanical mechanism as the *RBC passage* and the *Hb viscosity transitions*, with additional hydrophobic patches forming abruptly at T_W_ (*23*).T_W_ thus separates two different physical states of the Hb surface on the temperature scale (*20,37,68,126,127*) (Fig. 2A).

### Hemoglobin Crystal Thermal Transitions (Hb crystal transition)

Water is a ubiquitous component of protein crystals, typically amounting to 30—70 % of total crystal volume. Gevorkian et al. grew monoclinic crystals of human oxyhemoglobin and measured the dynamic Young modulus (modulus of elasticity) E as a function of temperature between 25 and 50°C (*128,129*). Intriguingly, it changed abruptly at several transition temperatures and remained constant in between (Fig. 3E).

**Figure 3E:**
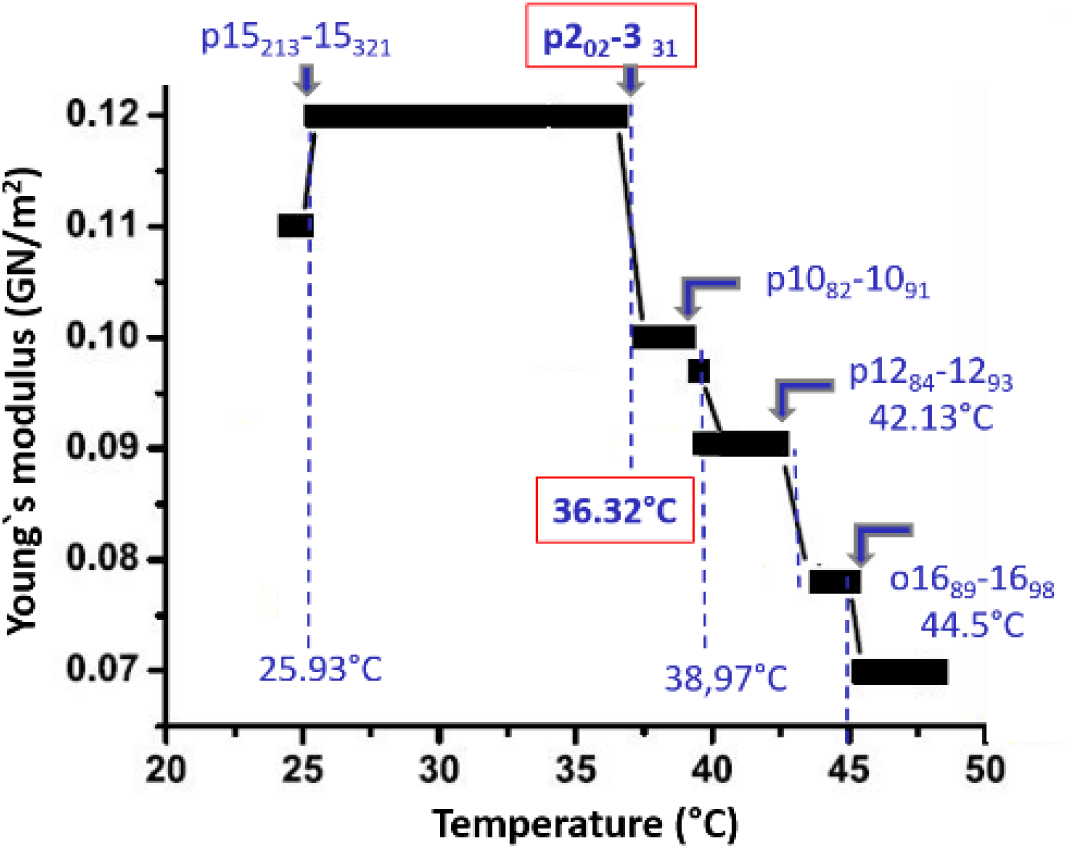
Hemoglobin crystal transition. (with kind permission (129)): Temperature profile of Young’s modulus, E, of human oxyhemoglobin crystals measured with an oscillating two-plate system (approx.. 5 kHz). E changes in sharp steps and remains constant in between. At T>49°C crystals clove to pieces. IWQ transitions and their characteristic temperatures TW (blue) are assigned to observed critical temperatures TC

The strength of lattice interactions, and hence the Young modulus, depends on the ambient humidity of the crystal, with E being approx. 8 times larger at 75% relative humidity than at 95%; the crystal becomes softer due to water absorption (*129*). Similar experiments have been reported for lysozyme crystals (*130*). It is interesting that the Young modulus increases abruptly at T_W_ = 25.93°C, while it decreases successively at all higher transition temperatures. Conceptually, a protein crystal can be regarded as an extremely concentrated protein solution, in which positions and orientations of individual molecules are restrained by lattice contacts while the majority of hydration water is preserved. We postulate that the increase in Young’s modulus at T_W_ = 25.93°C (p15_312_-15_213_) results from an abrupt change in the hydrogen bonding architecture at the protein-water interface and/or moderate alterations of the protein surface affecting crystal lattice contacts.

Obviously, structural alterations must be moderate enough to not disrupt the lattice contacts, especially since non-fixated crystals were used in these studies. An increased density of hydrogen bonds, or other non-covalent interactions, would explain the observation of the crystal becoming less elastic (*37,96,131*). The new binding state is maintained from T ≥ 25.93°C until changing to another H-bridge arrangement at T_W_ = 36.32°C (p2_02_-3_31_). At T_W_=36.32°C, the crystal suddenly “softens”. Note that the resonance method applied here for measuring the Young modulus generates shear forces between Hb molecules and adjacent water layers (*92*). We hypothesize that, analogous to the viscosity transition described above, an increase in the effective hydrophobicity of Hb due to rotational transitions of water in the protein-water interface leads to a decrease in the density of protein-water hydrogen bonds. This might enable the onset of low-friction sliding, rationalising the sudden drop of Young’s modulus.

In the range between T_W_ = 25.93 °C and T_W_ =36.32 °C further rotational transitions are expected, but these do not appear to have any effect on Young’s modulus, either because the energy input is too small to cause a change or because the alterations are silent w.r.t. crystal elasticity.

### Colloid osmotic pressure (COP) of RBCs in autologous plasma (Hb-COP transition)

In a series of in vitro experiments, we measured the COP of human RBCs resuspended in autologous plasma with an average RBC content of 77.6±5.3% by volume (RBC-in-plasma sample) versus autologous plasma alone (plasma sample) at 29°C ≤ T ≤ 39.5°C (see Fig. 4 in *23*). This was to elucidate as closely as possible the in vivo environment of the hemoglobin-water interaction inside RBCs (*23,120*). The COP of both samples increases linearly and parallelly, with the plasma sample COP consistently 0.27 kPa higher. It is reasonable to assume that this is due to a small proportion of plasma proteins adhering to the RBC surfaces in the RBC-plasma sample, thus reducing the free plasma protein concentration and it’s COP. While the COP of the plasma sample continued to increase linearly with temperature, the parallelism of the COPs ended at T_C_ = 37.1±0.2 °C. From here on, the COP of the RBC-plasma sample decreased rapidly, at 39.5°C being 0.73 kPa lower than that of the plasma sample. Our assumption is that from T_C_ upwards cytosolic water leaks from the RBC into the surrounding plasma. This dilution effect causes the plasma protein concentration to decrease and the total COP to decline in parallel. Based on the IWQ-Model, the transition at T_C_ induces partial hydrophobisation on the Hb surfaces, triggering cytosolic Hb aggregation and thus reducing the number of colloid osmotically active particles. As a result, cytosolic water is released into the extracellular space (*27,31,132–134*).

**Figure 4A:**
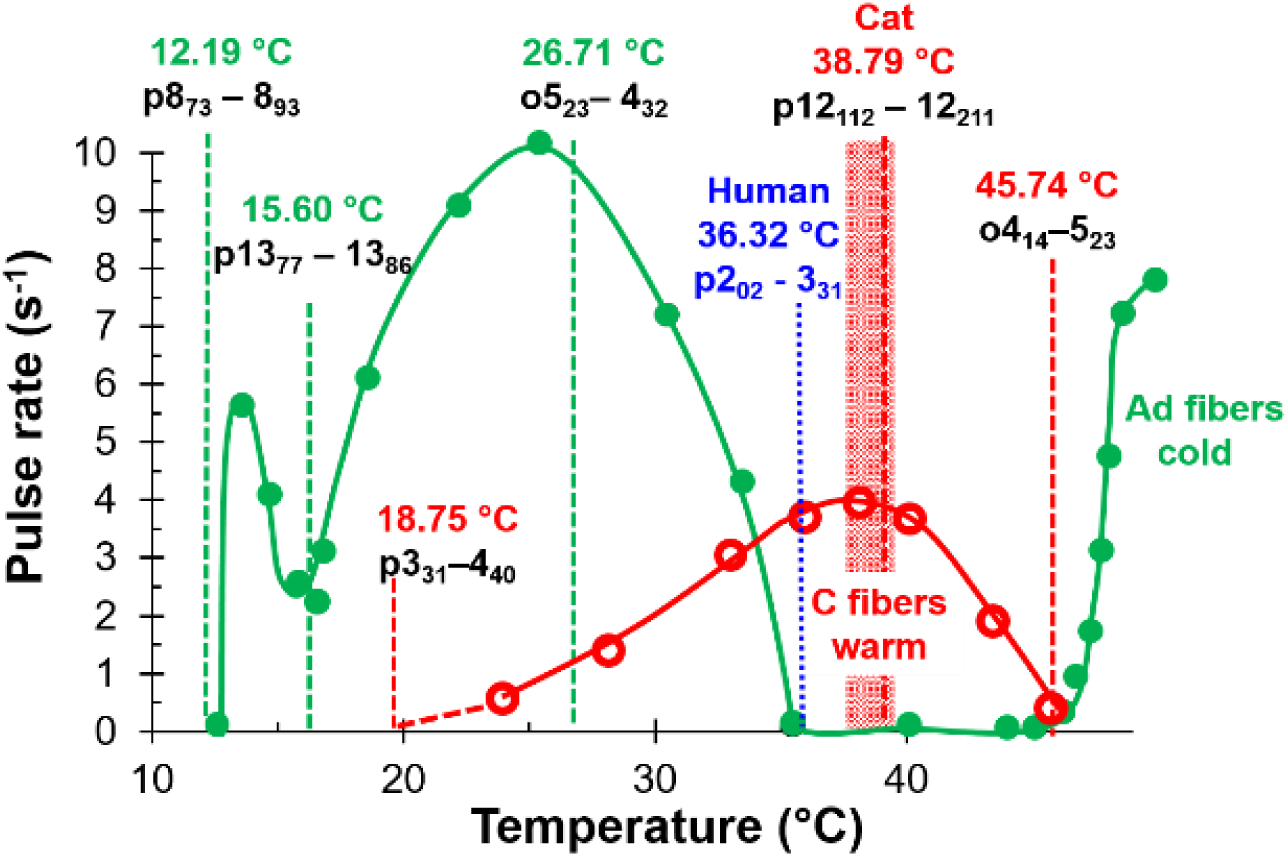
Temperature sensitive neurons (cat) (with kind permission (41)): Steady state firing rate of individual cold-sensitive Aδ−fibers (green) and warm-sensitive C fibers (red) after thermal stimulation of the lingual nerve of cats at constant temperatures between 10 and 50°C. The normal body temperature range of cats is shaded red. Temperatures at which the sensitivity of the two fiber types is turned on, turned off, or turned over are assigned IWQ reference temperatures.

The Hb-COP transition takes place in the full absence of fluid dynamic shear forces. Taking into account the above considerations and the technical shortcomings of T_C_ measurement, we hypothesize that the COP transition is another manifestation of IWQ transition p2_02_-3_31_ with characteristic temperature T_W_=36.32°C, proposed above as a reference for basal human body temperature T_B, human_. The release of water from the cytosol of RBCs may represent a contribution to homeostasis during fever in humans (*40,46,135*). From the COP data, it can be estimated that an adult with a fever temperature of 39.5°C transfers about 500 ml of water from RBCs into the blood plasma, partially compensating for the loss of sweat water (*23*). Analogously, we assume that additional water is recruited from muscle cells during fever due to aggregation of myoglobin setting on possibly as well at TC (*135,136*). A possible physiological relationship between the release of water by the RBCs from T_W_ = 36.32°C on and the increase in the rate of skin temperature-related sweating as the core body temperature exceeds approx. 36.5°C is only hinted at here (*4*).

## IWQ transitions of thermosensitive TRP transmembrane channels

### Neuron Thermal Sensitivity

Using the local temperature distribution in an organism for regulation of its body temperature requires sophisticated biological sensing mechanisms (*45,49,54,137*). First, a neuronal temperature transition on the organ level is discussed here (*42*). The study involved the stimulation of heat-sensitive splanchnic nerves in the dorsal wall of the abdominal cavity of rabbits through the perfusion of thermodes with water at temperatures ranging from 12°C to 50°C. Up to a critical temperature, Tc approx. 36-38°C, neural afferent activity remained constant. It then changes to an almost linear increase with temperature (see Fig. 3 in *42*). Applying the IWQ Model, the rectal body temperature of the rabbit TB approx. 38.4±0.6°C can be assigned the IWQ reference temperature T_W_=38.69°C (o12_012_-12_111_). Thus, the authors provide a (rare) example of a critical switching temperature at the organ level, T_C,organ_ (*39,42,61,139*). It seems reasonable to assume that such higher-order switching temperatures are intrinsically related to the critical temperatures, T_C_, of key molecular temperature sensors, in this case the heat-sensitive TRP ion channels. These include the noxious heat-sensitive TRPV1 (VR1) channel as well as the warm sensitive TRPV3 channel (*140–142*) (discussed below).

A further example is the temperature profile of the action potential frequency recorded from individual thermosensitive neuronal fibers, the cold fibers (Aδ-fibers) and the warm fibers (C fibers), after thermal stimulation of *cat* tongue nerves (Fig. 4A) (*14,31,41,44,145*).

The cold-sensitive Aδ-fibers clearly show multiple regions of different temperature sensitivity (*146*). The starting point of continuous discharge correlates with T_W_=12.19°C (p8_73_-8_93_). There is an intermediate minimum and a maximum located close to T_W_=15.6°C (p13_77_-13_86_) and T_W_=26.71°C (o4_23_-5_32_), respectively. The activity stops around T_W_=36.32°C (p2_02_-3_31_). Interestingly a second temperature sensitive range sets on at approx. T_W_=45.74°C (o4_14_-5_23_). We recognize an analogy to the “cold sensation paradox” that is typically observed in humans at temperatures above 45°C. The warm-sensitive C-fibers instead begin firing at approx. T_W_=18.75°C (p3_31_-4_40_), and reach a maximum rate close to T_W_=38.79°C (p12_112_-12_211_), which is within the cat’s normal body temperature range, T_B_ approx. 38.3-39.2°C. Sensitivity dissipates at T_W_=45.74°C (o4_14_-5_23_), the same temperature at which the cold-sensitive Aδ-fibers resume their temperature sensitivity (*147*). In generating afferent temperature signals over the entire temperature range, C- and Aδ-fibers complement each other (*139,143,148*).

### Temperature sensing TRP channels

The TRP channels TRPM8 (cold sensor), TRPV3 (warm sensor), and TRPV1 (noxious heat sensor) are molecular temperature sensors with exceptionally high Q_10_ values in certain temperature ranges (*14,43,45,47,50,51,54,57,58,66,94,137,141,144*).

The cold sensitive TRPM8 is expressed in approx. 15% of small dorsal root ganglion neurons and is activated below T_C_=25°C (Fig. 4B) (*146*). Whole-cell patch clamp data obtained with HEK293 cells showed robust temperature-activated currents.

**Figure 4B:**
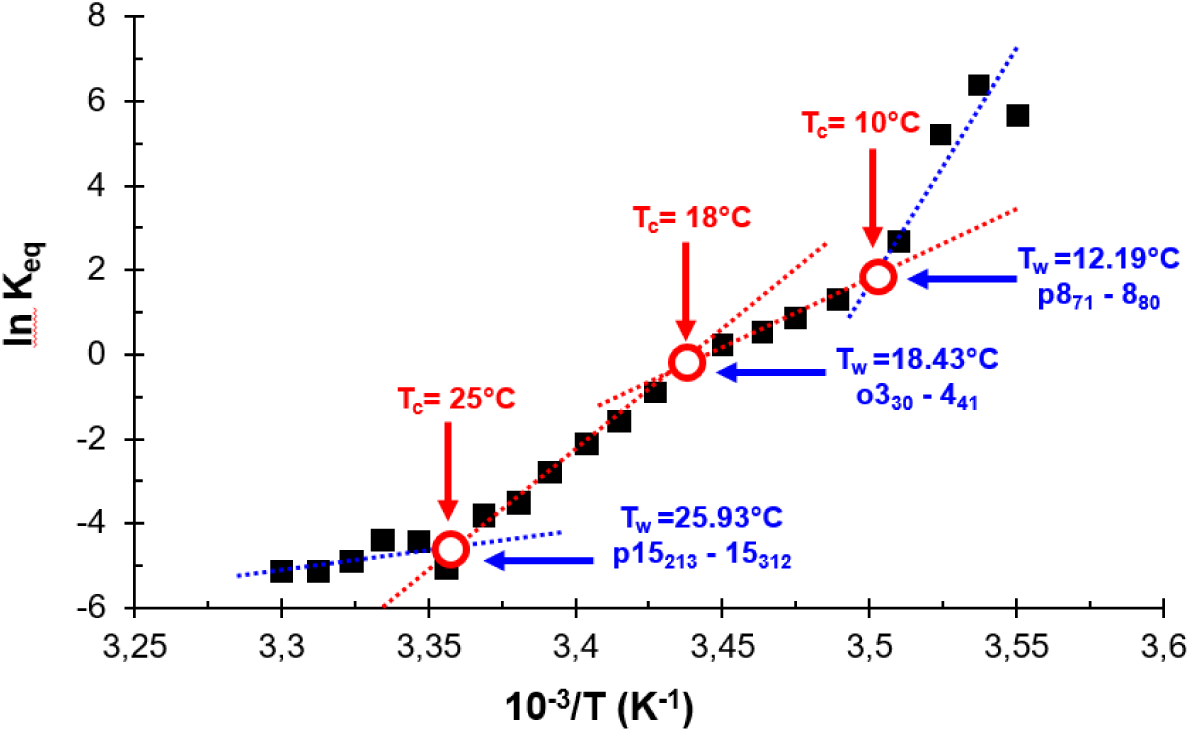
Cold sensitive TRPM8. (with kind permission (146), Figure 2C): Van’t Hoff plot of steady-state membrane channel activity, Keq, measured by whole-cell patch-clamping at 60 mV holding potential. The authors performed a linear regression analysis for two temperature regimes (red dotted lines), 10°C to 18°C (Q10=3.3) and 18°C to 25°C (Q10=23.8). However, we identify two more regimes, one at T<10°C and another shallow one at T>25°C (blue dotted lines). The three critical temperatures were assigned to IWQ reference temperatures. Deviations between TW vs. TC reflect experimental temperature uncertainties.

Channel activation is usually described using a two-state model, i.e., under the simplified assumption of a single open and closed state, respectively. Figure 4B displays a van’t Hoff plot of the equilibrium constant K_eq_= [open]/[closed]. The plot exhibits several regions of differential temperature dependence (Q_10_), with linear segments separated by inflection points at T_C_=10°C, 18°C, and 25°C. Assigned IWQ reference temperatures are indicated.

At T_C_=18°C the enthalpy and entropy of channel opening (related to slope and intercept of the linear fit) change by ΔΔH=52 kcal mol^−1^ and ΔΔS=174 cal mol^−1^ K^−1^. Based on the IWQ model, the energy required for the underlying structural perturbation is covered by rotational transitions of water at T_W_ = 18.43°C (o3_30_-4_41_). At this T_W,_ one water molecule absorbs an energy of 0.58 kcal/mol (wavenumber of 202.69 cm^−1^). The change in gating enthalpy at T_C_=18°C corresponds to about 90 rotational transitions of water molecules per tetramer (equations (1) and (2)). Since the channel is composed of four identical subunits, the changes to protein conformation should be statistically symmetric, and further synchronization may occur via allosteric coupling. For TRPM8, two additional enthalpy jumps were observed at T_C_=25°C and T_C_=10°C. Above 25°C, the TRPM8 channel follows a shallower phase at very low activity. Below T_C_=10°C, the authors described the regime as ‘saturated’, but the data may indicate a low-temperature range with a very high Q_10_. The TRPM8 channel therefore shows a different sensitivity in several temperature ranges, which is also typical for cold-sensitive A*δ* fibers (Fig. 4A).

The warm sensitive TRPV3 is predominantly expressed in skin keratinocytes but also in the tongue, dorsal root ganglia, spinal cord, and brain (*48,54,55,150*).

Figure 4C shows a Van’t Hoff plot obtained from the equilibrium constant, K_eq_, of wild-type mouse TRPV3 reconstituted into lipid Nano discs. Above a critical temperature at approx. 36 °C, the equilibrium constant increases rapidly with temperature (Q_10_ = 27.0 ± 7.4). The limited number of data points did not allow the T_C_ to be determined with high accuracy. For the same reason, the existence of further critical temperatures below 36°C cannot be ruled out. Nevertheless, the assumption T_C_ approx. T_B,mouse_ approx. T_W_=36,32°C (p2_02_-3_31_) appears reasonable. This transition is accompanied by changes in gating enthalpy and entropy of ΔΔH = 91.2 ± 3.6 kcal mol^−1^ and ΔΔS = 287.5 ± 11.7 cal mol^−1^ K^−1^, respectively (*54*). Rotational transitions, p2_02_-3_31_ (215.13 cm^−1^) at Tw=36.32°C, absorb an energy of 0.62 kal mol^−1^. According to equation (1) and (2) a sum of 147 transitions of free water molecules per tetramer (about 37 per monomer) would be equivalent to enable this enthalpy jump.

**Figure 4C:**
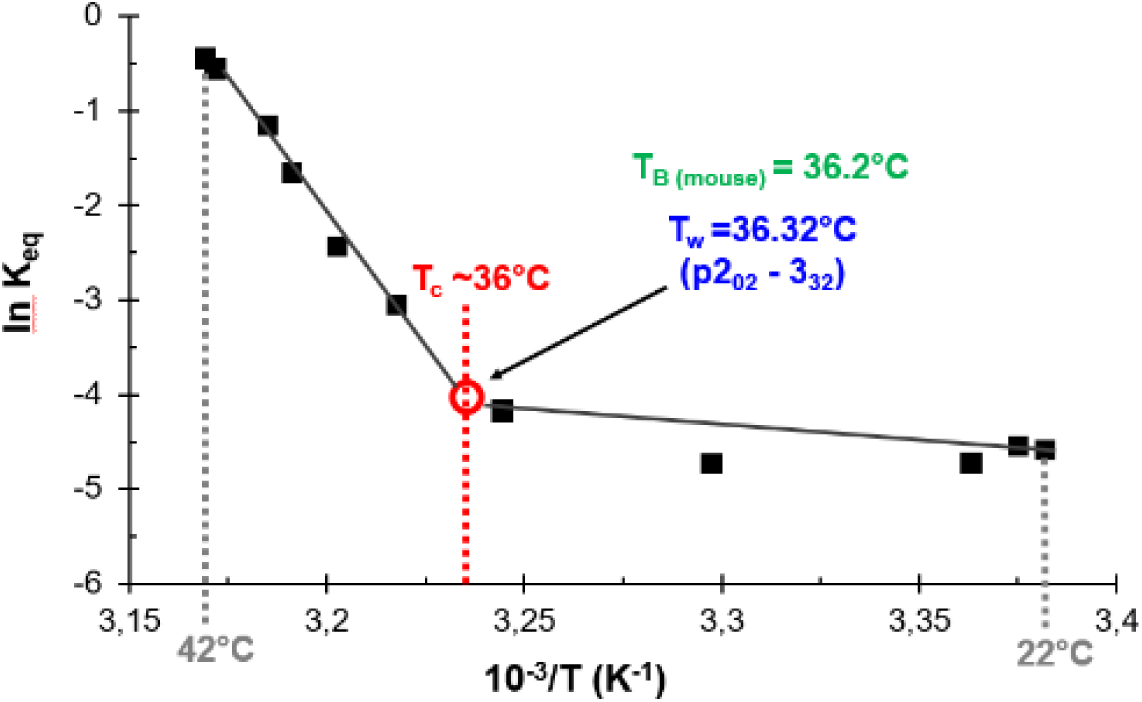
Warm sensitive TRPV3 (with kind permission (54), extended data Fig. 1C): TRPV3 is thermosensitive in the range of temperatures between TRPM8 and TRPV1. The Van’t Hoff plot of the equilibrium constant measured by patch-clamping reveals two Q10 regimes separated by a kink at TC∼36°C corresponding to mouse body temperature TB, mouse=36,2°C and TW=36,32°C (p202-331).

In their cryo-EM studies of TRPV3 reconstituted in Nano discs, Nadezhdin et al. (*54*) identified two distinct responses to heat depending on the type of Nano disc used, one corresponding to the canonical open state and a one in an intermediate conformational state which they interpret as sensitized (activated but still closed). Since their heat activation protocol involves oscillation between 25°C and 42°C, crossing the major kink in temperature regimes discussed above, it is tempting to speculate that their sensitized state may share features with the high-Q_10_ state defined by the van’t Hoff plot. Notably, the closed-to-sensitized transition involves secondary structure changes and rigid-body motions together affecting a large portion of the molecule, which seems consistent with the distributed nature of a solvent-mediated effect, while the actual channel opening transition was more localized. Obviously, cryo-EM would only be able to resolve temperature-dependent structural changes relaxing slowly compared to the cryocooling kinetics.

The TRPV1 channel (VR1) is sensitive to noxious heat (Fig. 4D) (*56,113,151*). Rat TRPV1 channels were expressed in oocytes as described by Catarina et al. and single channel activity was measured by patch-clamping (*49,57*).

**Figure 4D:**
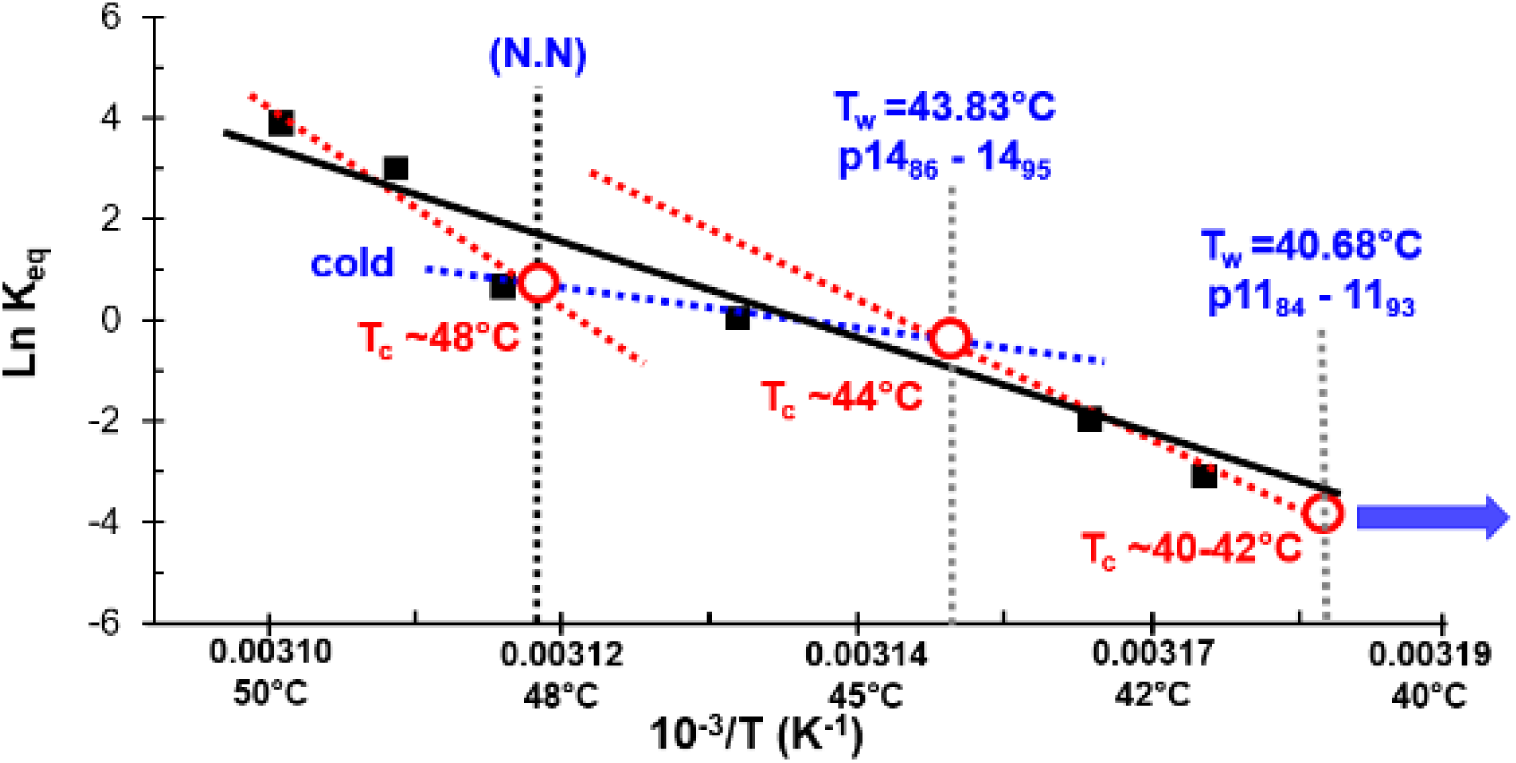
Noxious heat sensitive TRPV1. (with kind permission (151)): Shown is a Van’t Hoff plot generated from the equilibrium constant K_eq_ for rat TRPV1 channels expressed in oocytes. Single-channel inside-out patch clamp recordings were taken. Individual TRPV1 channel activity started between T_C_∼40-42°C with T_W_=40,68°C (p11_84_-11_93_). Although the authors performed a simple linear regression, our detailed analysis revealed two additional critical temperatures, T_C_=44°C (T_W_=43.83°C, p14_86_-14_96_) and TC=48°C (T_W_=N.N.), and four Q10 regimes.

The authors’ linear regression between 40°C and 50°C (black line) yielded a gating enthalpy of 150±13 kcal/mol (*151*). As with the VRPM8 channels, it seems possible to identify more than one temperature regime. Accordingly, the channel activity switches between higher enthalpy (red dashed lines) and lower enthalpy Q_10_ regimes (blue dashed line).

### IWQ Mechanism of Temperature-Induced Switching of TRP Channel activity

The molecular mechanisms responsible for thermally activating TRP channels remain to be fully elucidated, but available data indicate that it involves conformational changes in different regions of the 3D structure as well as allosteric interactions (Nadezhdin et al., 2021) (Fig. 4E). Thermodynamic considerations suggest that, assuming only a single open-closed transition, channel gating should be temperature dependent, favouring similar conformational changes for both heat and cold stimulation. This is due to the temperature dependence of the heat capacity change at constant pressure ΔCp modulating ΔH and ΔS and hence K_eq_ of the opening transition (*50*). An increase in heat capacity upon channel opening was proposed to be caused by exposure of about 50 nonpolar hydrophobic side chains (10-20 per TRP protomer), representing less than 2% of all side chains in typical TRP channels. However, the experimental data discussed above (Fig. 4B-D) indicate the presence of several states featuring individual temperature sensitivities, which correspond to slightly different closed and open structures. Within each Q_10_ interval, the Clapham and Miller model should apply in principle (*50*). However, the Van’t Hoff plot’s approximate linearity and the constancy of both ΔH and ΔS within these intervals suggest that the influence of ΔCp is minimal at best. In contrast, abrupt changes in specific heat are likely associated with transitions between different Q_10_ regimes (Fig. 4F). Such an increase in ΔCp is consistent with our hypothesis that IWQ transitions at discrete temperatures, T_W_ can lead to protein conformational changes involving an increase in (effective) hydrophobicity. Provided that these changes are fully reversible, the switching enthalpy is released upon return to the previous temperature regime (*152*).

**Figure 4E:**
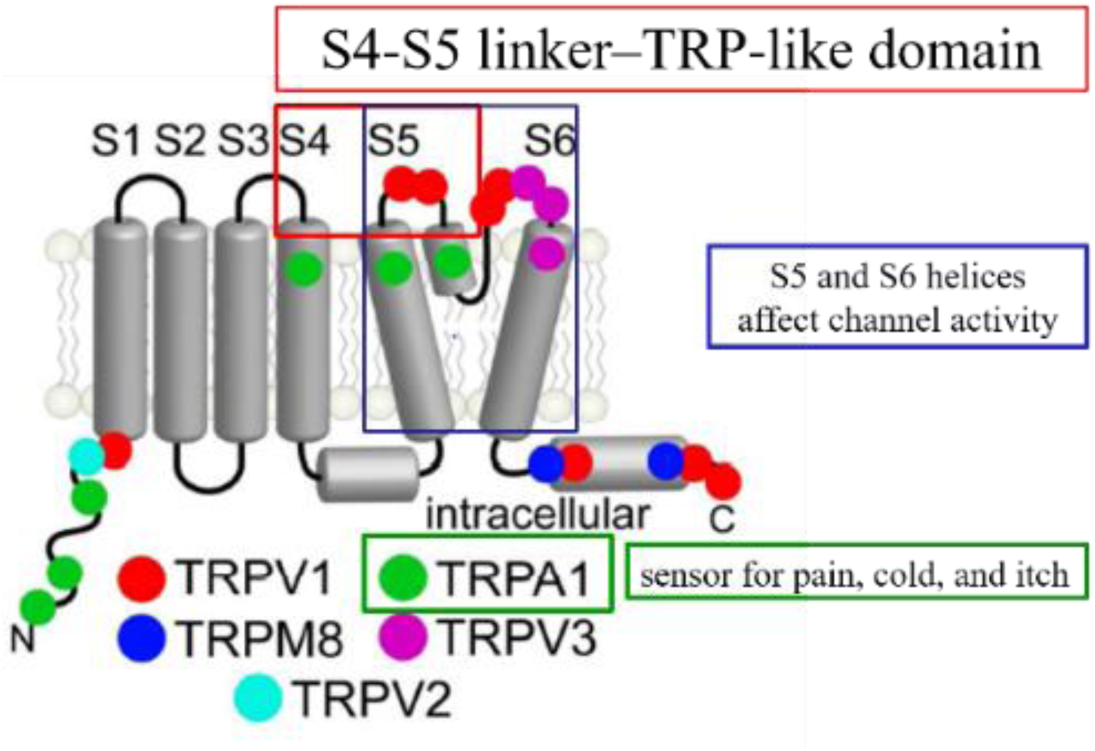
Regions identified as thermo-active in different TRP channels. (with kind permission Hilton et al. 2015 (58), figure 4). They are mostly located in solvent-accessible portions of the protein, facing either the extracellular space or the cytosol, which is in agreement with principal IWQ model requirements.

The free energies associated with conformational changes in TRP channels are typically on the order of 100 kcal/mol, which is equivalent to approx. 160 water molecules absorbing 0.62 kcal/mol at the rotational transition p2_02_-3_31_ (*146*). Even if this is only a rough estimate, two conclusions can be drawn: firstly, the excitation of a single free water molecule in the interface involves a non-negligible amount of energy, even when viewed on a macromolecular scale. Second, among the thousands of water molecules comprising the first hydration layer of a large protein molecule, a small percentage undergoing quantum absorption will be sufficient to elicit a biological effect.

The water accessible portions of TRP channels are in contact with either the extracellular medium or the cytosol (Fig. 4E) (*57,58,66*).

With respect to the IWQ model, we propose that the thermally susceptible areas are located here because this is where the enthalpies required for switching can be acquired via water quantum transitions. The primary (localized) structural changes originating from energy uptake can spread as a “conformational wave” throughout the protein, similar to the closed-to-sensitized transition observed by Nadezhdin et al. (*54*).

Supported by allosteric coupling, these alterations will affect the entire tetrameric channel, including its transmembrane region and the gating mechanism, finally resulting in the switch between thermal regimes observed experimentally (Fig. 4F).

**Figure 4F:**
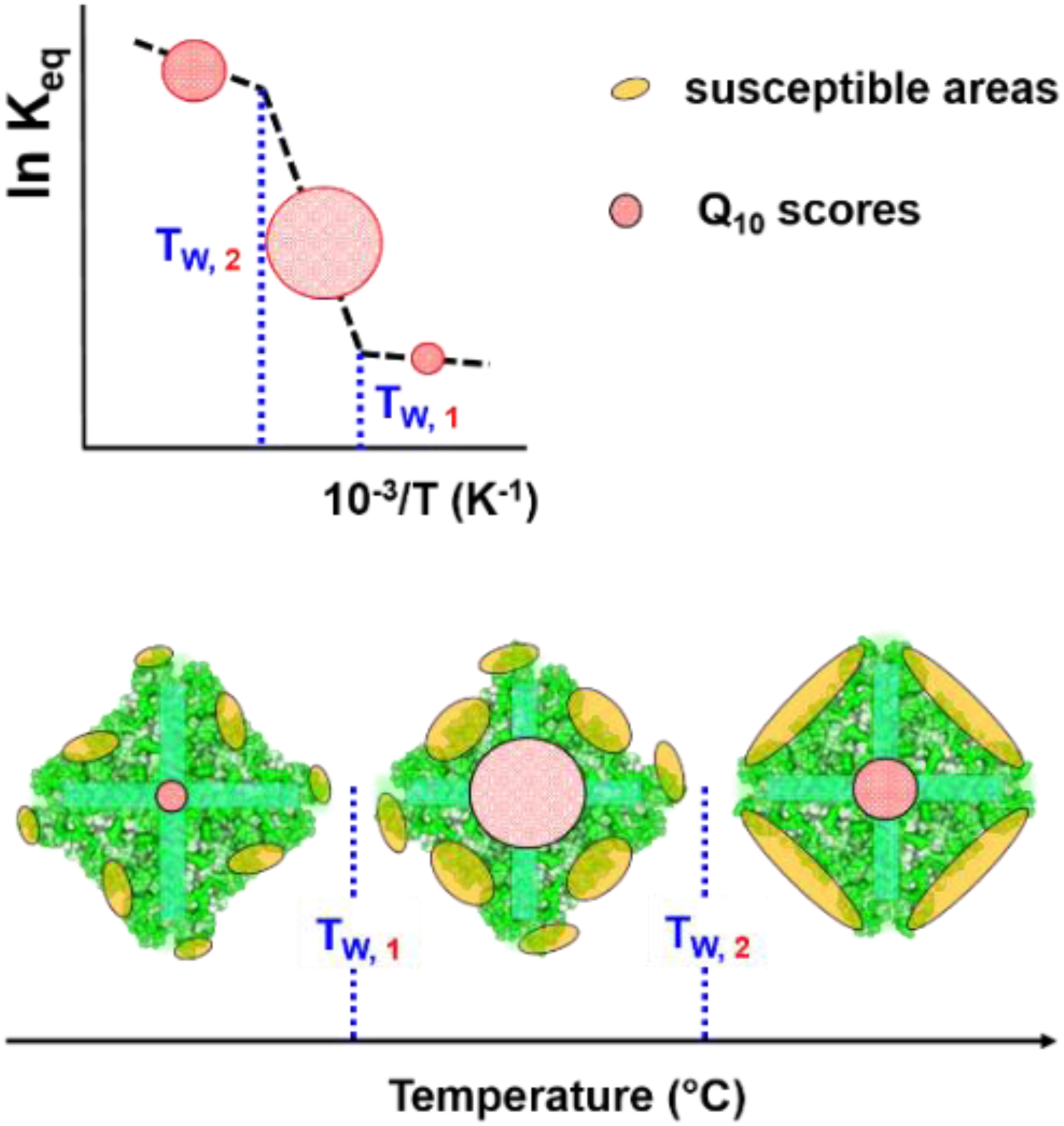
**Below:** This schematic representation shows a top view of a homotetrameric TRP channel (green) (compare to Hilton et al.(66)) with susceptible areas (light brown) for different Q_10_ intervals. The size and surface distribution of these areas are protein-specific and change discontinuously at the IWQ temperatures T_W,1_ and T_W,2_ (blue), concurrent with switches between Q_10_ regimes. The size of the central circle illustrates the temperature sensitivity, i.e. the Q_10_ value between the IWQ temperatures. **Above:** This schematic van’t Hoff plot demonstrates typical TRP channel patch clamp data with IWQ temperatures T_W_ indicated. At each transition temperature, energy is absorbed locally in the susceptible areas. The allosteric interaction of these regions finally enables the switching to another Q_10_ regime.

The precise localization of the thermosensitive domains is still uncertain, and experimental studies have yielded conflicting results. Intriguingly, even a large-scale mutagenesis effort involving thousands of random exchanges in rat TRPV1 has failed to clearly map thermal susceptibility to a defined domain or region of the protein (*80*). The data did, however, indicate that mutations reducing hydrophobicity tended to reduce thermosensitivity but not the response to ligands.

The authors interpreted their observations in terms of the heat capacity hypothesis proposed by Clapham and Miller (*50*). As pointed out above, this model is based on single open and closed states and, in contrast to the IWQ-Model, cannot account for the switches between distinct temperature regimes featuring constant gating enthalpies and entropies. In fact, both the seemingly distributed nature of the effect and the dependence on apolar side chains are consistent with the postulate of the IWQ Model that energy uptake at defined temperatures can be mediated by rotational activation of free water molecules. While we expect the distribution of thermo-susceptible areas in the protein-water interface to be complex, it is certainly not random. Any given IWQ model based rotational transition will result in a modified surface topology with an altered distribution of hot spots for generation of free water molecules, which are available for energy uptake as a different resonance temperature is reached (Fig. 4F). Obviously, the information governing the thermal response of TRP channels, and hence the differences among channel types and paralogues, must ultimately be encoded in the amino acid sequence.

## IWQ Model implications for thermal homeostasis

### Body Temperatures of Homeothermic Species and the IWQ temperatures

Being homeothermic organisms, birds and mammals benefit from a strict regulation of their body temperature (T_B_) for constant performance and responsiveness. T_B_ is optimized during evolution, and significant deviations from normal values can be fatal. Poikilothermic species, on the other hand, tolerate changes in their core temperatures over a wide range. The process of thermoregulation itself has been studied extensively for decades and is not further considered here (*1,2,3,7,44,153*). Our key question is: How does a homeothermic organism “know” what the basal value of its core body temperature is, regardless of all external and internal influences (*10*).

Figure 5 shows histograms of experimentally determined body temperatures for 596 mammals and 632 birds. Intriguingly, we find that both the main peaks and several secondary peaks can be assigned IWQ temperatures. In general, body temperatures are not normally distributed (Jarque Bera Test), and the frequency distribution for birds features stronger negative skew and higher kurtosis than that of mammals. The fine structure of both distributions may be due to real variation, but probably also reflects technical problems such as unequal sampling of taxa and rounding effects. Nevertheless, it is interesting to note that, despite the significant skewness, the peak value of the mammalian distribution, T_B, peak, mammals_=36,45°C, determined after smoothing closely matches the arithmetic mean, T_B, mammals_=36.46°C, both being close to the commonly-quoted standard human body temperature, T_B,human_=36.6°C (*10*). T_B, peak, mammals_ as well as T_B, human_, in turn, correlate with the specific water rotational transition temperature T_W_=36.32 °C (p2_02_->3_31_), as postulated by the IWQ model. The maximum of the frequency distribution of birds is found at T_B, peak, birds_= 42.2°C, and the respective IWQ temperature is T_W_=42.13°C (p12_84_-12_93_). The Eastern Bluebird (Sialia sialis) represents a typical avian species with a body temperature of 42.2°C.

**Figure 5:**
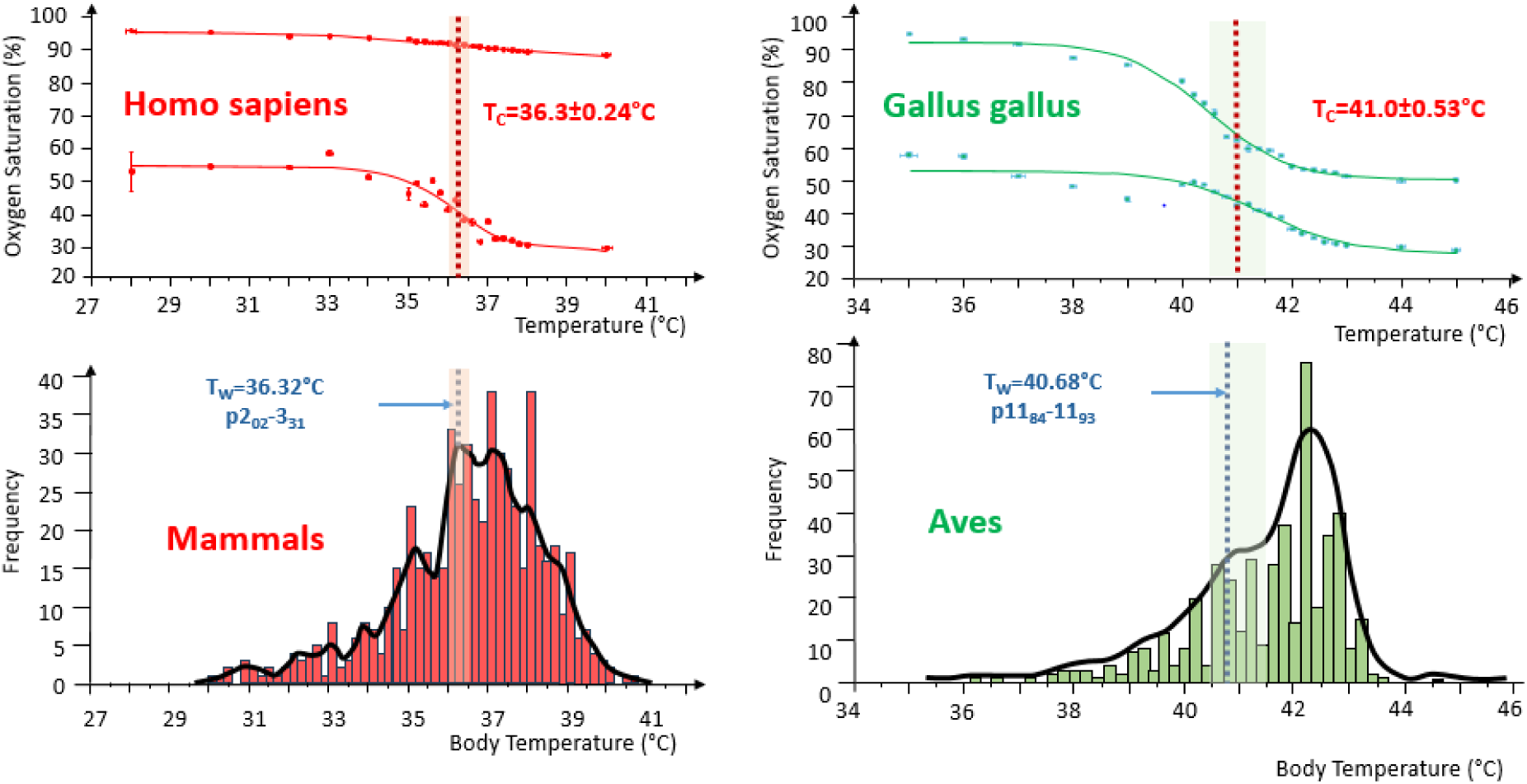
Oxygen saturation transition: **Bottom row**: Frequency distributions of body temperatures of mammals (N=596, TB,mean =36.46±1.88°C and birds (N = 464, TB,mean=41.45±1.29°C) (mean±SD) 154 with the envelope curve showing the major and minor maxima. **Upper row:** Temperature dependence of the hemoglobin oxygen saturation (left, human; right, chicken) at atmospheric oxygen partial pressure (upper curves) and 50% oxygen partial pressure (below). A sigmoidal relation was discovered in all cases. The turning points, T_C_, correlate remarkably well with distinct rotational transition temperatures, T_W_, within the limits of the measuring errors (dashed areas). The data represent mean±1SD of oxygen saturation for each temperature (N=3).

### Hemoglobin function and body temperature – the Oxygen saturation transition

Using a variety of methods, we have demonstrated correlations between critical temperatures in the behaviour of intact RBCs or Hb solutions and the body temperatures of the respective species (*23*). By comparing the hemoglobin sequences of these species, conserved and divergent sequence segments were identified as indicated in Zerlin et al. (*24,25*). Hemoglobin molecular regions with higher sequence divergence tend to be closer to the surface and involve both α and β subunits of the hemoglobin tetramer, including the α-β interfaces. We hypothesize that both the overall globin fold and the alpha-beta architecture of hemoglobin contain information about body temperature (*71,75,155*). Although the oxygen-binding myoglobin is a monomer, it is conceivable that it might feature a similar species-specific temperature sensitivity as observed for hemoglobin.

For homiotherme species, binding of oxygen to hemoglobin during the allosteric T-R transition is a vital function (*156*). It is therefore plausible that, during the evolution of organisms, body temperatures were selected to optimally support oxygen transport by hemoglobin in the respective species. During oxygen binding, hemoglobin undergoes a reversible change in its three-dimensional structure from a tense state (T-state) with lower affinity to a relaxed state (R-state) featuring high affinity, due to an allosteric cooperative effect. In general, oxygen affinity of hemoglobin is well-known to decrease with increasing temperature, corresponding to a shift of the oxygen binding isotherm towards higher partial pressures (*157,158*).

We reasoned that the temperature-dependent transitions in structure and properties observed for hemoglobin or intact RBCs may also manifest in oxygen transport. To verify this prediction, we re-assessed the temperature dependence of oxygen saturation (SO_2_) for both mammalian (human) and avian (chicken) hemoglobin at two constant partial pressures pO_2_, (1) using ambient air, i.e., 150 mmHg of oxygen, and (2) 3.6% oxygen in nitrogen, corresponding to 27 mmHg of oxygen, the half-saturation pressure for human hemoglobin at human body temperature). Indeed, these recent experiments revealed sigmoidal relations. For human hemoglobin at ambient pO_2_, this was a weak trend, with an almost linear decrease in oxygen saturation of 0.67%/K (r=0.96). Around half-saturation partial oxygen pressure, however, we observed a pronounced sigmoidal decrease (maximum slope of –8.4 %/K) in human hemoglobin oxygen saturation with temperature (Fig. A5, left, above). Moreover, the inflection point of this curve, T_C_= 36.3±0,24°C, was identical to the IWQ transition temperature, T_W_=36.32°C (p2_02_-3_31_) and thus close to the basal value of the human body temperature, T_B,Human_=36.6°C (*10*). To our surprise, we found that for chicken hemoglobin oxygen saturation exhibited a highly significant sigmoidal decrease already at ambient oxygen partial pressure, with T_C_= 41.0 ± 0,53°C and a slope at T_C_ of −14.3%/K, whereas at 27 mmHg of oxygen, the slope at T_C_ (−7.7%/K) was smaller (Fig. A5, right, above). Consistent with our observations with the human protein, the T_C_ for chicken hemoglobin was, within the limits of measurement accuracy, identical to the chicken body temperature, T_B, chicken_ (40.6-41.7°C). This value corresponds to the IWQ transition temperature T_W_=40.68°C (p11_84_–11_93_).

Apparently, the oxygen binding isotherms do not just shift smoothly with temperature but display a particularly strong temperature dependence (appearing as a transition at constant partial pressure) around critical temperatures T_C_. To the best of our knowledge, this oxygen saturation transition has not been described before. Based on these observations, we expect similar results for other homeothermic species. If such a correlation was confirmed, this would support the generalised conjecture T_C, species_=T_B, Species_=T_W, species_, with T_W_ serving as IWQ reference temperature for the respective species.

### IWQ Model Mechanism of the Hb-Oxygen Saturation transition

The process of binding oxygen in the R state is exothermic, while releasing oxygen in the T state is endothermic, explaining the general decrease of oxygen affinity with increasing temperature (*158*). If the temperature exceeds T_C_ and the Hb oxygen saturation drops as observed, then energy must be supplied to the Hb molecule in order to increase the T-state probability, causing a fraction of bound oxygen to be released (Fig. 5). Considering the IWQ-Model, the energy required is invested to the Hb-water Interface in a specific and sudden manner. The critical temperature, T_C_, for the oxygen saturation transition of human Hb can be assigned to the same rotational quantum transition of water (p2_02_-3_31_, with T_W_=36.32°C) as the *Hb crystal transition* and the *RBC passage transition* discussed above. These temperature transitions of very different physical parameters exhibit similar steepness, supporting the view that they share the same cause.

In analogy to the TRP channels (Fig. 4F), the IWQ energy is stored in the susceptible areas existing in R-state hemoglobin around T_C_. The resulting alterations in the tertiary structures of α and β chains affect their elementary oxygen binding constants and/or their cooperativity, which ultimately leads to the distinct sigmoidal shape of the hemoglobin saturation curve around T_C_. In physiological terms, this is reminiscent of the Bohr effect (decrease of oxygen affinity upon lowering the pH) or the action of other inorganic (chloride) or organic modulators (2,3-bisphosphoglycerate). The difference is that rotational transitions alter the T-R equilibrium in response to a physical parameter rather than the chemical environment, and the temperature-dependent switch may or may not resemble the chemically induced ones. Regardless of structural details, the IWQ effect is likely to favor oxygen release in peripheral tissues at elevated temperatures, e.g. in skeletal muscle during exercise or under fever conditions. While the temperature sensitivity of human Hb oxygen saturation is low at atmospheric pO_2_, or the pressure encountered in lung capillaries (100 mmHg), it increases with rising temperature to as much as 8.4 %/K at T_C_ if pO_2_ is close to the half-saturation level (Fig. 5). The overall result not only appears consistent with physiological demands but also is expected to usually synergize with the Bohr Effect.

Another physiological effect is associated with the ‘work of solvation’ (WoS) required during the transition from the T-state to the R-state (*76,85*). It must be invested in the (colder) lung alveoli during Hb oxygenation in order to bring water molecules to the Hb surface exposed during the transition from the T-state to the R-state. According to Colombo et al., water molecules act as allosteric effectors during HB oxygenation (T->R transition). In the course of this conformational change, human hemoglobin requires approximately 60 water molecules to cover the additional surface area (420-600 Å^2^, roughly 4% of the sovent-accessible surface) that is exposed in the relaxed state. This implies an osmotic WoS in the physiological medium of 0.2 kcal/mol. The WoS is released upon return to the T state during deoxygenation in peripheral tissues, making the process appear as a physiological heat pump. Since this process is coupled to oxygen delivery, it should share the same temperature dependence and hence be pronounced when the temperature exceeds T_C_. Furthermore, part of the water absorbed during oxygen binding is released again as saturation drops at T≥T_C_.

While monomeric myoglobin does not exhibit an T-R transition, it may still be subject to an oxygen saturation transition triggered by IWQ effects. Indeed, we were able to confirm a structural transition similar to hemoglobin via CD spectroscopy; the implications for oxygen delivery to muscle tissue are uncertain at present.

## Summary and Conclusions

This article presents a model for understanding discontinuities in the thermal behaviour of proteins at discrete transition temperatures, with a focus on homeothermic species, in particular humans. The basic assumption is: At critical temperatures, thermal energy is absorbed locally in the protein-water interface, potentially resulting in conformational changes and an associated switch in the protein’s function. The surface distribution of this energy uptake is specific to the protein (and the species) as well as the transition temperature considered. As the underlying mechanism, we propose rotational quantum transitions of temporarily free water molecules in the protein-water interface (*62,65,87,109*). This hypothesis is supported by the established rotational excitation of free water molecules in fullerene cages and the experimental demonstration of such transitions in aqueous protein and DNA solutions by means of four-photon spectroscopy (*91,94,99,103*).

The IWQ model does not make any assumptions about specific structural properties of biomolecules but relies solely on the temperature dependence of the protein-water interface, including changes to the hydrogen bonding architecture and the topography of hydrophobic regions. Since the proposed mechanism represents a generic route for the water-mediated transfer of enthalpy to all kinds of co-solvents or solutes, downstream processes might be modulated in every physically possible way: in addition to favoring higher-energy states of macromolecules, as assumed in the discussion above, energy might, e.g., be utilized to overcome activation barriers, enabling transitions which are otherwise kinetically hindered though thermodynamically favorable.

This model now enables us to explain the transitions observed decades ago for human RBCs as well as solutions of human hemoglobin, including the *RBC passage transition*, the *Hb viscosity transition*, and the *CD secondary structure transition*: in all cases the inflection point is virtually indistinguishable from the resonance temperature assigned to quantum transition p2_02_-3_31_ (T_W_=36.32°C) of *para*-water. Comparison across a number of mammalian and avian hemoglobins suggests that the switching temperature strongly correlates with the species’ body temperature and must be evolutionarily engraved into the structure of the protein.

The IWQ model implies that pronounced changes in structure and properties as a function of temperature should not be limited to soluble proteins but may equally affect all sorts of membrane-bound proteins and ultimately any biomolecule, with potentially profound impact on all aspects of life.

As an instructive example, we reviewed available patch clamp data on temperature-sensitive TRP channels. Unfortunately, published information on equilibrium constants in the temperature range considered here is limited. While previous models have rationalized the general temperature-dependence of gating, the remarkable switching between distinct thermal regimes at well-defined temperatures has remained enigmatic. Similar to the case of hemoglobin, the IWQ model provides a solution to the discontinuity problem by assuming that thermal energy is absorbed with particularly high efficiency at temperatures supporting rotational quantum transitions of water. This way, each type of channel can utilize its own repertoire of transition temperatures, allowing for subtype and species specificity; alterations in the amino acid sequence can widely modulate the thermal response but not the position of the candidate transition temperatures because the latter are dictated by quantum physics. Finally, based on the examples shown, we emphasize the enormous potential of patch-clamp experiments for assessing the temperature dependence of channel activity. Particular attention should be paid to the temperature accuracy directly at the cell patch.

The initial question of the paper was: Is there a physically defined reference for body temperature? We propose that it is set by the *hemoglobin oxygen saturation transition*. This hypothesis is supported by our recent investigation of human and chicken Hb oxygen saturation as a function of temperature. Both recordings feature a sigmoidal relation (Fig. 5, top row), with inflection points coinciding with other hemoglobin transitions as well as with the body temperatures of the respective species. Based to these observations, temperature qualifies as another allosteric modulator causing a prominent shift in the T-R equilibrium (and hence the oxygen saturation isotherm), which is centered on basal body temperatures. The quantum transition temperature T_W_=36.32°C of the rotational transition p2_02_-3_31_ therefore suggests itself as the physical reference value of *human* body temperature. We expect that such an assignment of body temperatures to specific rotational water transitions is also possible for other homeothermic species, but this needs to be confirmed by future investigations.

Owing to the underlying physical mechanism, there is no reason to assume that IWQ transitions would not occur in the proteins of poikilothermic species. Since these have no well-defined body temperature, it is conceivable that their essential proteins are relatively stable and active over a wide range of temperatures. Nevertheless, a correlation of critical temperatures affecting certain body functions with IWQ transition temperatures can be expected. For example, the ambient temperature to which turtle eggs are exposed during hatching determines the sex of the offspring which means that temperature controls gene expression in these animals (*159,160*). Obviously, IWQ transitions might affect transcription or translation by “switching” nucleic acid-binding proteins, and previous experiments have revealed water quantum rotation also in the presence of naked DNA (*99,161,162*). Similar considerations apply to temperature-dependent effects in all organisms, examples being microbial metabolism and plant growth (*163,164*).

Regardless of the specifics of the system under study, the sharp and often unanticipated transitions in certain temperature-dependent biological processes suggest several critical requirements for future experiments: first and foremost, an accuracy of < ±0.2°C in assessing temperature inside the sample volume studied, and a sufficiently long equilibration time after technically reaching a temperature level. Furthermore, we recommend experiments with human cells and proteins or DNA at “body temperature” to be carried out in steps of 0.2°C around T_W_ = 36.32°C. Results obtained “at laboratory temperature” have their place in history and are of limited value.

In summary, we think that temperatures playing a decisive role in physiology, including the basal body temperature in homeothermic species, can be traced back to rotational quantum transitions of *ortho-* or *para-*water. The IWQ model provides a novel framework for understanding previously unexplained experimental observations. Casting the concept into a concise formula: Proteins sense and water sets critical physiological temperatures.

## Materials and Methods

### Oxygen saturation measurements of hemoglobin

*Fresh human blood was collected via finger prick from a healthy donor into heparin-coated capillaries (BRAND, Wertheim, Germany). Brown nick chicken blood conserved in Alsever’s solution was acquired from Labor Dr. Merk (Ochsenhausen, Germany). Blood samples were centrifuged with isotonic phosphate saline and plasma was discarded. The packed red cells were hemolysed and further diluted with 0.1 M phosphate buffer, pH 7.0* (*165,166*)*, and centrifuged to remove the stroma. The diluted hemoglobin samples were directly used for spectrophotometric analysis. The UV-VIS spectrophotometer V-550 (JASCO, Japan) was coupled with a circulation thermostat PT80 (Krüss, Hamburg, Germany) which heats both sample and reference chambers. Phosphate buffer was used as blank and reference. A 3D printed cuvette cover that accommodates the temperature probe TDIP15 (Pyroscience, Aachen, Germany), O2 sensor OXF1100-OI (Pyroscience, Aachen, Germany) and holes for gas inlet and overpressure outlet was used. For measurements involving oxyhemoglobin, the samples were analyzed in ambient air. To create 50% oxygenated hemoglobin, the specimens were purged with a gas mixture consisting of 3.6% oxygen and the remainder nitrogen (Linde, Pullach, Germany). The oxygen saturation was calculated using Pittman et al.’s method* (*167*)*. The oxygen saturation data were fitted to sigmoid curves to determine the inflection points using MatLab*.

## Acknowledgments

We would like to thank all the scientists, including our students, who have accompanied us on this journey and helped us to make it a success. We would also like to express our gratitude to you, esteemed colleagues in the scientific community, for your invaluable contributions as role models, educators, colleagues, and employees. Please accept this expression of our sincerest gratitude for your invaluable assistance.

## Author contributions

Conceptualization: GMA, OW, AA

Methodology: GMA, ISF

Investigation: GMA, AA, ISF

Visualization: SD, GMA

Supervision: AA, GMA

Writing – original draft: GMA, OW

Writing – review & editing: OW, AA, SD, ISF

## Data and materials availability

All data and further information are available on request from the corresponding author.

## References and Notes

[1] C. A. Wunderlich, On the temperature in diseases : a manual of medical thermometry, 1st ed. London: The New Sydenham society, 1871.

[2] H. T. Hammel and J. B. Pierce, “Regulation of Internal Body Temperature,” Annu. Rev. Physiol., vol. 30, no. 1, pp. 641–710, Mar. 1968, doi: 10.1146/annurev.ph.30.030168.003233.

[3] C. M. Blatteis, “A personal recollection: 60 years in thermoregulation,” Temperature, vol. 3, no. 1, Jan. 2016, doi: 10.1080/23328940.2016.1148524.

[4] C. H. Wyndham, A. R. Atkins, “A physiological scheme and mathematical model of temperature regulation in man,” Pflugers Arch. Eur. J. Physiol., vol. 303, no. 1, 1968, doi: 10.1007/BF00586824.

[5] C. Jessen, “Thermal afferents in the control of body temperature,” Pharmacol. Ther., vol. 28, no. 1, Jan. 1985, doi: 10.1016/0163-7258(85)90085-3.

[6] C. L. Tan and Z. A. Knight, “Regulation of Body Temperature by the Nervous System,” Neuron, vol. 98, no. 1, Apr. 2018, doi: 10.1016/j.neuron.2018.02.022.

[7] N. R. Cabej, “Control Systems and Determination of Phenotypic Traits in Metazoans,” in Epigenetic Principles of Evolution, Elsevier, 2019.

[8] M. O. Moreira, Y. Qu, and J. J. Wiens, “Large-scale evolution of body temperatures in land vertebrates,” Evol. Lett., vol. 5, no. 5, pp. 484–494, Oct. 2021, doi: 10.1002/evl3.249.

[9] M. Protsiv, C. Ley, J. Lankester, T. Hastie, and J. Parsonnet, “Decreasing human body temperature in the United States since the Industrial Revolution,” Elife, vol. 9, Jan. 2020, doi: 10.7554/eLife.49555.

[10] Z. Obermeyer, J. K. Samra, and S. Mullainathan, “Individual differences in normal body temperature: longitudinal big data analysis of patient records,” BMJ, vol. 1, p. j5468, Dec. 2017, doi: 10.1136/bmj.j5468.

[11] D. D. McKemy, “Temperature sensing across species,” Pflugers Archiv European Journal of Physiology, vol. 454, no. 5. pp. 777–791, Aug. 2007, doi: 10.1007/s00424-006-0199-6.

[12] M. G. Artmann, H. Schmid-Schoenbein, “Erythrocyte in a capillary junction (Video),” https://www.youtube.com/watch?v=_Jlu8Df-VKc, 1989.

[13] G. M. Artmann, C. Kelemen, D. Porst, G. Büldt, and S. Chien, “Temperature Transitions of Protein Properties in Human Red Blood Cells,” Biophys. J., vol. 75, no. 6, Dec. 1998, doi: 10.1016/S0006-3495(98)77759-8.

[14] G. M. Artmann, “Erythrocyte aspiration into a 1.3µm pipette at 35°C (video),” https://www.youtube.com/watch?v=bzYBU8oNZ-Q, 1994.

[15] Artmann G, “Erythrocyte aspiration into a 1.3µm pipette at 37°C (video),” 1994. https://www.youtube.com/watch?v=s3xA47T81uA (accessed Jun. 01, 2024).

[16] O. K. Baskurt, A. Temiz, and H. J. Meiselman, “Original Contribution effect of superoxide anions on red blood cell rheologic properties,” 1998.

[17] S. Weinbaum, Y.-S. J. Li, and G. W. Schmid-Schönbein, “A Lifetime Achievement in Bioengineering: Professor Shu Chien,” Ann. Biomed. Eng., vol. 47, no. 11, Nov. 2019, doi: 10.1007/s10439-019-02390-2.

[18] T. M. Fischer, “On the energy dissipation in a tank-treading human red blood cell,” Biophys. J., vol. 32, no. 2, pp. 863–868, 1980, doi: 10.1016/S0006-3495(80)85022-3.

[19] R. M. Hochmuth, “Micropipette aspiration of living cells,” J. Biomech., vol. 33, no. 1, Jan. 2000, doi: 10.1016/S0021-9290(99)00175-X.

[20] Y.-B. Yan, Q. Wang, H.-W. He, and H.-M. Zhou, “Protein Thermal Aggregation Involves Distinct Regions: Sequential Events in the Heat-Induced Unfolding and Aggregation of Hemoglobin,” Biophys. J., vol. 86, no. 3, pp. 1682–1690, Mar. 2004, doi: 10.1016/S0006-3495(04)74237-X.

[21] G. M. Artmann et al., “Circular dichroism spectra of human hemoglobin reveal a reversible structural transition at body temperature,” Eur. Biophys. J., vol. 33, no. 6, Oct. 2004, doi: 10.1007/s00249-004-0401-8.

[22] C. Kelemen, “Analyse des phasensprungartigen Einsetzens der Passage humaner Erythrozyten durch Mikropipetten bei kritischen Temperaturen.,” Laboratory for Cell Biophysics, (G.M. Artmann), FH Aachen & Institute für Biologische Informationsverarbeitung (G. Bueldt), Juelich, 1999.

[23] G. M. Artmann et al., “Hemoglobin senses body temperature,” Eur. Biophys. J., vol. 38, no. 5, Jun. 2009, doi: 10.1007/s00249-009-0410-8.

[24] I. Digel et al., “Body Temperature-Related Structural Transitions of Monotremal and Human Hemoglobin,” Biophys. J., vol. 91, no. 8, Oct. 2006, doi: 10.1529/biophysj.106.087809.

[25] K. F. T. Zerlin et al., “Structural transition temperature of hemoglobins correlates with species’ body temperature,” Eur. Biophys. J., vol. 37, no. 1, Nov. 2007, doi: 10.1007/s00249-007-0144-4.

[26] A. M. Stadler et al., “From Powder to Solution: Hydration Dependence of Human Hemoglobin Dynamics Correlated to Body Temperature,” Biophys. J., vol. 96, no. 12, Jun. 2009, doi: 10.1016/j.bpj.2009.03.043.

[27] A. M. Stadler et al., “Cytoplasmic Water and Hydration Layer Dynamics in Human Red Blood Cells,” J. Am. Chem. Soc., vol. 130, no. 50, Dec. 2008, doi: 10.1021/ja807691j.

[28] A. M. Stadler, L. van Eijck, F. Demmel, and G. Artmann, “Macromolecular dynamics in red blood cells investigated using neutron spectroscopy,” J. R. Soc. Interface, vol. 8, no. 57, Apr. 2011, doi: 10.1098/rsif.2010.0306.

[29] C. Kelemen, S. Chien, and G. M. Artmann, “Temperature Transition of Human Hemoglobin at Body Temperature: Effects of Calcium,” Biophys. J., vol. 80, no. 6, Jun. 2001, doi: 10.1016/S0006-3495(01)76232-7.

[30] P. Cioni, G. B. Strambini, “Effect of heavy water on protein flexibility,” Biophys. J., vol. 82, no. 6, pp. 3246–3253, 2002, doi: 10.1016/S0006-3495(02)75666-X.

[31] A. Stadler, “La Dynamique Moléculaire dans les Globules Rouges,” Université Joseph Fourier – Grenoble 1, Grenoble, 2014.

[32] A. Clarke and P. Rothery, “Scaling of body temperature in mammals and birds,” Funct. Ecol., vol. 0, no. 0, pp. 071029083929001-???, Oct. 2007, doi: 10.1111/j.1365-2435.2007.01341.x.

[33] A. Clarke, P. Rothery, and N. J. B. Isaac, “Scaling of basal metabolic rate with body mass and temperature in mammals,” J. Anim. Ecol., vol. 79, no. 3, pp. 610–619, May 2010, doi: 10.1111/j.1365-2656.2010.01672.x.

[34] H. S. Frank et al., “REPRODUCTIVE BIOLOGY,” 2012.

[35] R. A. Eagle et al., “Isotopic ordering in eggshells reflects body temperatures and suggests differing thermophysiology in two Cretaceous dinosaurs,” Nat. Commun., vol. 6, no. 1, p. 8296, Nov. 2015, doi: 10.1038/ncomms9296.

[36] A. M. Stadler et al., “Thermal fluctuations of haemoglobin from different species: adaptation to temperature via conformational dynamics,” J. R. Soc. Interface, vol. 9, no. 76, Nov. 2012, doi: 10.1098/rsif.2012.0364.

[37] N. B. Rego and A. J. Patel, “Understanding Hydrophobic Effects: Insights from Water Density Fluctuations,” Annu. Rev. Condens. Matter Phys. is, vol. 13, pp. 303–327, 2022, doi: 10.1146/annurev-conmatphys.

[38] G. I. Makhatadze, P. L. Privalovt, “Heat Capacity of Proteins I. Partial Molar Heat Capacity of Individual Amino Acid Residues in Aqueous Solution: Hydration Effect,” 1990.

[39] P. L. Privalov and G. I. Makhatadze, “Heat Capacity of Proteins II. Partial Molar Heat Capacity of the Unfolded Polypeptide Chain of Proteins: Protein Unfollding Effects,” 1990.

[40] S. F. Morrison, K. Nakamura, “Central Mechanisms for Thermoregulation,” Annu. Rev. Physiol., vol. 12, p. 57, 2018, doi: 10.1146/annurev-physiol-020518.

[41] Y. Zotterman, “Special Senses: Thermal Receptors,” Annu. Rev. Physiol., vol. 15, no. 1, pp. 357– 372, Mar. 1953, doi: 10.1146/annurev.ph.15.030153.002041.

[42] W. Riedel, G. Siaplauras, and E. Simon, “Intra-abdominal thermosensitivity in the rabbit as compared with spinal thermosensitivity,” Pflügers Arch. Eur. J. Physiol., vol. 340, no. 1, pp. 59–70, 1973, doi: 10.1007/BF00592197.

[43] F. Yeh, A. Jara-Oseguera, and R. W. Aldrich, “Implications of a temperature-dependent heat capacity for temperature-gated ion channels,” Proc. Natl. Acad. Sci. U. S. A., vol. 120, no. 24, 2023, doi: 10.1073/pnas.2301528120.

[44] E. A. Tansey, C. D. Johnson, and C. D. Johnson, “Staying Current Recent advances in thermoregulation,” Adv Physiol Educ, vol. 39, pp. 139–148, 2015, doi: 10.1152/advan.00126.2014.-Ther.

[45] J. A. Lamas, L. Rueda-Ruzafa, and S. Herrera-Pérez, “Ion Channels and Thermosensitivity: TRP, TREK, or Both?,” International journal of molecular sciences, vol. 20, no. 10. NLM (Medline), May 14, 2019, doi: 10.3390/ijms20102371.

[46] K. Song et al., “The TRPM2 channel is a hypothalamic heat sensor that limits fever and can drive hypothermia,” Science (80-.)., vol. 353, no. 6306, pp. 1393–1398, Sep. 2016, doi: 10.1126/science.aaf7537.

[47] D. Julius, “TRP channels and pain,” Annual Review of Cell and Developmental Biology, vol. 29. pp. 355–384, Oct. 2013, doi: 10.1146/annurev-cellbio-101011-155833.

[48] D. E. Clapham, “TRP channels as cellular sensors,” 2003. [Online]. Available: www.nature.com/nature.

[49] M. J. Caterina, M. A. Schumacher, M. Tominaga, T. A. Rosen, J. D. Levine, and D. Julius, “The capsaicin receptor: a heat-activated ion channel in the pain pathway,” 1997.

[50] D. E. Clapham and C. Miller, “A thermodynamic framework for understanding temperature sensing by transient receptor potential (TRP) channels,” Proc. Natl. Acad. Sci. U. S. A., vol. 108, no. 49, pp. 19492–19497, Dec. 2011, doi: 10.1073/pnas.1117485108.

[51] D. M. Bautista et al., “The menthol receptor TRPM8 is the principal detector of environmental cold,” Nature, vol. 448, no. 7150, Jul. 2007, doi: 10.1038/nature05910.

[52] S. S. Rosenbaum T, “TRP Ion Channel Function in Sensory Transduction and Cellular Signaling Cascades,” in TRP Ion Channel Function in Sensory Transduction and Cellular Signaling Cascades., W. Liedtke and S. Heller, Eds. Boca Raton (FL): CRC Press/Taylor & Francis, 2007.

[53] S. K. Mishra, S. M. Tisel, P. Orestes, S. K. Bhangoo, and M. A. Hoon, “TRPV1-lineage neurons are required for thermal sensation,” EMBO J., vol. 30, no. 3, Feb. 2011, doi: 10.1038/emboj.2010.325.

[54] K. D. Nadezhdin et al., “Structural mechanism of heat-induced opening of a temperature-sensitive TRP channel,” Nat. Struct. Mol. Biol., vol. 28, no. 7, pp. 564–572, Jul. 2021, doi: 10.1038/s41594-021-00615-4.

[55] A. K. Singh, L. L. McGoldrick, L. Demirkhanyan, M. Leslie, E. Zakharian, and A. I. Sobolevsky, “Structural basis of temperature sensation by the TRP channel TRPV3,” Nat. Struct. Mol. Biol., vol. 26, no. 11, pp. 994–998, Nov. 2019, doi: 10.1038/s41594-019-0318-7.

[56] D. D. Luu, A. M. Owens, M. D. Mebrat, and W. D. Van Horn, “A molecular perspective on identifying TRPV1 thermosensitive regions and disentangling polymodal activation,” Temperature, vol. 10, no. 1, pp. 67–101, 2023, doi: 10.1080/23328940.2021.1983354.

[57] M. Kim et al., “Evidence that the TRPV1 S1-S4 membrane domain contributes to thermosensing,” Nat. Commun., vol. 11, no. 1, Dec. 2020, doi: 10.1038/s41467-020-18026-2.

[58] J. K. Hilton, P. Rath, C. V. M. Helsell, O. Beckstein, and W. D. Van Horn, “Understanding thermosensitive transient receptor potential channels as versatile polymodal cellular sensors,” Biochemistry, vol. 54, no. 15, pp. 2401–2413, Apr. 2015, doi: 10.1021/acs.biochem.5b00071.

[59] E. Schrödinger, What is Life, 1st ed. Camebridge: Cambridge University Press, 1944.

[60] J. Crovisier et al., “The Spectrum of Comet Hale-Bopp (C/1995 O1) Observed with the Infrared Space Observatory at 2.9 Astronomical Units from the Sun,” Science (80-.)., vol. 275, no. 5308, pp. 1904–1907, Mar. 1997, doi: 10.1126/science.275.5308.1904.

[61] A. F. Bunkin, G. A. Lyakhov, A. A. Nurmatov, and A. V. Rezov, “Four-photon spectroscopy of collective interactions between water molecules and electromagnetic radiation,” Phys. Rev. B, vol. 52, no. 13, pp. 9360–9363, 1995, doi: 10.1103/PhysRevB.52.9360.

[62] P. L. Chapovsky and L. J. F. Hermans, “NUCLEAR SPIN CONVERSION IN POLYATOMIC MOLECULES,” 1999. [Online]. Available: www.annualreviews.org.

[63] A. F. Bunkin, V. I. Grachev, G. A. Lyakhov, and A. A. Nurmatov, “Four-photon polarization spectroscopy of water in a millimeter-wave radiation field,” 1998.

[64] H. J. Bakker, “Water’s response to the fear of water,” Nature, vol. 491, no. 7425, Nov. 2012, doi: 10.1038/491533a.

[65] A. F. Bunkin and S. M. Pershin, “Observation of water isotopes and spin-isomers rotational transitions induced by four-wave mixing in liquid,” in Journal of Raman Spectroscopy, 2008, vol. 39, no. 6 SPEC. ISS., pp. 726–729, doi: 10.1002/jrs.1992.

[66] J. K. Hilton, M. Kim, and W. D. Van Horn, “Structural and Evolutionary Insights Point to Allosteric Regulation of TRP Ion Channels,” Acc. Chem. Res., vol. 52, no. 6, pp. 1643–1652, Jun. 2019, doi: 10.1021/acs.accounts.9b00075.

[67] P. Agre, “Aquaporin Water Channels,” Biosci. Rep., vol. 24, no. 3, Jun. 2004, doi: 10.1007/s10540-005-2577-2.

[68] M. Meuwly and M. Karplus, “The Functional Role of the Hemoglobin-Water Interface,” Oct. 2021, [Online]. Available: http://arxiv.org/abs/2110.03201.

[69] G. H. Pollack, I. L. Cameron, and D. N. Wheatley, “Water and the cell,” Water Cell, no. September, pp. 1–354, 2006, doi: 10.1007/1-4020-4927-7.

[70] V. Adrian Parsegian, “Protein-water interactions,” 2002.

[71] S. K. Pal and A. H. Zewail, “Dynamics of water in biological recognition,” Chem. Rev., vol. 104, no. 4, pp. 2099–2123, Apr. 2004, doi: 10.1021/cr020689l.

[72] K. A. Dill, “Theory for the folding and stability of globular proteins,” Biochemistry, vol. 24, no. 6, pp. 1501–1509, Mar. 1985, doi: 10.1021/bi00327a032.

[73] H. Kendrew, J.C.; Bodo, G., Dintzis, H.M., Parrish, R.G., and Wyckoff, “A Three-dimensional Model of the Myoglobin Molecule Obtained by X-ray Analysis,” Nature, vol. 181, pp. 662–666, 1958, doi: 10.1038/181662a0.

[74] A. F. Cullis, H. M. Dintzis, and M. F. Perutz, “X-ray analysis of haemoglobin,” 1958.

[75] H. Frauenfelder et al., “Thermal Expansion of a Protein,” Biochemistry, vol. 26, pp. 254–261, 1987, [Online]. Available: 10.1021/bi00375a035.

[76] M. Colombo, D. Rau, and V. Parsegian, “Protein solvation in allosteric regulation: a water effect on hemoglobin,” Science (80-.)., vol. 256, no. 5057, May 1992, doi: 10.1126/science.1585178.

[77] A. D. Friesen and D. V. Matyushov, “Local polarity excess at the interface of water with a nonpolar solute,” Chem. Phys. Lett., vol. 511, no. 4–6, pp. 256–261, Aug. 2011, doi: 10.1016/j.cplett.2011.06.031.

[78] J. G. Davis, K. P. Gierszal, P. Wang, and D. Ben-Amotz, “Water structural transformation at molecular hydrophobic interfaces,” Nature, vol. 491, no. 7425, pp. 582–585, Nov. 2012, doi: 10.1038/nature11570.

[79] S. Pezzotti et al., “Spectroscopic Fingerprints of Cavity Formation and Solute Insertion as a Measure of Hydration Entropic Loss and Enthalpic Gain,” Angew. Chemie - Int. Ed., vol. 61, no. 29, Jul. 2022, doi: 10.1002/anie.202203893.

[80] J. O. Sosa-Pagán, E. S. Iversen, and J. Grandl, “TRPV1 temperature activation is specifically sensitive to strong decreases in amino acid hydrophobicity,” Sci. Rep., vol. 7, no. 1, Dec. 2017, doi: 10.1038/s41598-017-00636-4.

[81] F. Mallamace et al., “Energy landscape in protein folding and unfolding,” Proc. Natl. Acad. Sci. U. S. A., vol. 113, no. 12, pp. 3159–3163, Mar. 2016, doi: 10.1073/pnas.1524864113.

[82] E. Xi et al., “Hydrophobicity of proteins and nanostructured solutes is governed by topographical and chemical context,” Proc. Natl. Acad. Sci. U. S. A., vol. 114, no. 51, pp. 13345–13350, Dec. 2017, doi: 10.1073/pnas.1700092114.

[83] J. Liu, X. He, and J. Z. H. Zhang, “Structure of liquid water – a dynamical mixture of tetrahedral and ‘ring-and-chain’ like structures,” Phys. Chem. Chem. Phys., vol. 19, no. 19, pp. 11931–11936, 2017, doi: 10.1039/C7CP00667E.

[84] S. N. Timasheff, “Protein hydration, thermodynamic binding, and preferential hydration,” Biochemistry, vol. 41, no. 46, pp. 13473–13482, Nov. 2002, doi: 10.1021/bi020316e.

[85] D. Bulone, P. L. San Biagio, M. B. Palma-Vittorelli, and M. U. Palma, “The role of water in hemoglobin function and stability,” Science, vol. 259, no. 5099. pp. 1335–1336, 1993, doi: 10.1126/science.8446903.

[86] L. S. Rothman et al., “The HITRAN 2004 molecular spectroscopic database,” J. Quant. Spectrosc. Radiat. Transf., vol. 96, no. 2 SPEC. ISS., pp. 139–204, Dec. 2005, doi: 10.1016/j.jqsrt.2004.10.008.

[87] L. S. Rothman et al., “The HITRAN 2008 molecular spectroscopic database,” J. Quant. Spectrosc. Radiat. Transf., vol. 110, no. 9–10, pp. 533–572, Jun. 2009, doi: 10.1016/j.jqsrt.2009.02.013.

[88] V. I. Tikhonov and A. A. Volkov, “Separation of Water into Its Ortho and Para Isomers.” [Online]. Available: www.sciencemag.org/cgi/content/full/296/5577/2363/.

[89] A. Farkas, “Orthohydrogen, Parahydrogen and Heavy Hydrogen,” Nature, vol. 135, no. 3416, Apr. 1935, doi: 10.1038/135601a0.

[90] A. Kilaj, H. Gao, D. Rösch, U. Rivero, J. Küpper, and S. Willitsch, “Observation of different reactivities of para and ortho-water towards trapped diazenylium ions,” Nat. Commun., vol. 9, no. 1, p. 2096, Dec. 2018, doi: 10.1038/s41467-018-04483-3.

[91] A. F. Bunkin, “Low-frequency spectroscopy of four-photon scattering in aqueous solutions of biopolymers,” Biophys. (Russian Fed., vol. 57, no. 6, pp. 709–715, 2012, doi: 10.1134/S0006350912060036.

[92] L. D. Landau and E. M. Lifshitz, Theory of Elasticity, 3rd ed., vol. 7. Amsterdam…Tokyo: Elsevier Ltd., 1986.

[93] G. Buntkowsky et al., “Mechanisms of Dipolar Ortho/Para-H 2 O Conversion in Ice,” Z. Phys. Chem, vol. 222, pp. 1049–1063, 2008, doi: 10.1524.zpch.2008.5359.

[94] C. Beduz et al., “Quantum rotation of *ortho* and *para*-water encapsulated in a fullerene cage,” Proc. Natl. Acad. Sci., vol. 109, no. 32, pp. 12894–12898, Aug. 2012, doi: 10.1073/pnas.1210790109.

[95] S. Mamone et al., “Nuclear spin conversion of water inside fullerene cages detected by low-temperature nuclear magnetic resonance,” J. Chem. Phys., vol. 140, no. 19, May 2014, doi: 10.1063/1.4873343.

[96] F. L. Thiemann, C. Schran, P. Rowe, E. A. Müller, and A. Michaelides, “Water Flow in Single-Wall Nanotubes: Oxygen Makes It Slip, Hydrogen Makes It Stick,” ACS Nano, vol. 16, no. 7, pp. 10775– 10782, Jul. 2022, doi: 10.1021/acsnano.2c02784.

[97] M. Frunzi et al., “A photochemical on-off switch for tuning the equilibrium mixture of H 2 nuclear spin isomers as a function of temperature,” J. Am. Chem. Soc., vol. 133, no. 36, pp. 14232–14235, Sep. 2011, doi: 10.1021/ja206383n.

[98] H.-H. Limbach et al., “Novel Insights into the Mechanism of the Ortho/Para Spin Conversion of Hydrogen Pairs: Implications for Catalysis and Interstellar Water,” ChemPhysChem, vol. 7, no. 3, pp. 551–554, Mar. 2006, doi: 10.1002/cphc.200500559.

[99] A. F. Bunkin, A. A. Nurmatov, S. M. Pershin, R. S. Khusainova, and S. A. Potekhin, “Four-photon microwave laser spectroscopy of aqueous solutions of biopolymers,” Quantum Electron., vol. 37, no. 10, pp. 941–945, Oct. 2007, doi: 10.1070/qe2007v037n10abeh013609.

[100] P. Kautny et al., “Charge-transfer states in triazole linked donor-acceptor materials: Strong effects of chemical modification and solvation,” Phys. Chem. Chem. Phys., vol. 19, no. 27, pp. 18055– 18067, 2017, doi: 10.1039/c7cp01664f.

[101] J. Liu, X. He, and J. Z. H. Zhang, “Structure of liquid water-a dynamical mixture of tetrahedral and ‘ring-and-chain’ like structures,” Phys. Chem. Chem. Phys., vol. 19, no. 19, pp. 11931–11936, 2017, doi: 10.1039/c7cp00667e.

[102] A. F. Bunkin, G. A. Lyakhov, A. A. Nurmatov, and N. V Suyazov, “Applied Physics B Lasers and Optics Origin of low-frequency spectrum structure of four-photon scattering in liquid water,” 1998.

[103] A. F. Bunkin, S. M. Pershin, R. S. Khusainova, and S. A. Potekhin, “Spin isomeric selectivity of water molecules upon DNA hydration,” Biophysics (Oxf)., vol. 54, no. 3, Jun. 2009, doi: 10.1134/S0006350909030026.

[104] W. Gawlik, J. Kowalski, F. Trager, and M. Vollmer, “PHYSICAL REVIEW LETTERS New Mechanism for the Production of Optical Resonances with Subnatural Linewidths,” 1982.

[105] M. F. Andersen et al., “Quantized rotation of atoms from photons with orbital angular momentum,” Phys. Rev. Lett., vol. 97, no. 17, 2006, doi: 10.1103/PhysRevLett.97.170406.

[106] M. Maiuri, M. Garavelli, and G. Cerullo, “Ultrafast Spectroscopy: State of the Art and Open Challenges,” J. Am. Chem. Soc., vol. 142, no. 1, pp. 3–15, Jan. 2020, doi: 10.1021/jacs.9b10533.

[107] G. Fleming, Chemical applications of ultrafast spectroscopy, 1st ed. OSTI.GOV_6037887, 1986.

[108] S. M. Pershin, A. F. Bunkin, and V. L. Golo, “H2O monomers in channels of icelike water structures,” J. Exp. Theor. Phys., vol. 115, no. 6, Dec. 2012, doi: 10.1134/S1063776112130109.

[109] S. M. Pershin, “OPTICAL SPECTROSCOPY Coincidence of Rotational Energy of H 2 O Ortho-Para Molecules and Translation Energy near Specific Temperatures in Water and Ice,” Phys. Wave Phenom., vol. 16, no. 1, pp. 15–25, 2008, doi: 10.3103/S1541308X08010032.

[110] S. M. Pershin, “Conversion of ortho-para H2O isomers in water and a jump in erythrocyte fluidity through a microcapillary at a temperature of 36.6±0.3°C,” Phys. Wave Phenom., vol. 17, no. 4, Dec. 2009, doi: 10.3103/S1541308X09040025.

[111] N. V. Prabhu, K. A. Sharp, “Heat capacity in proteins,” Annu. Rev. Phys. Chem., vol. 56, pp. 521– 548, 2005, doi: 10.1146/annurev.physchem.56.092503.141202.

[112] V. L. Arcus et al., “On the Temperature Dependence of Enzyme-Catalyzed Rates,” Biochemistry, vol. 55, no. 12. American Chemical Society, pp. 1681–1688, Mar. 29, 2016, doi: 10.1021/acs.biochem.5b01094.

[113] A. Mugo, R. Chou, F. Chin, B. Liu, Q. X. Jiang, and F. Qin, “A suicidal mechanism for the exquisite temperature sensitivity of TRPV1,” Proc. Natl. Acad. Sci. U. S. A., vol. 120, no. 36, 2023, doi: 10.1073/pnas.2300305120.

[114] A. Sánchez-Moreno et al., “Irreversible temperature gating in trpv1 sheds light on channel activation,” 2018, doi: 10.7554/eLife.36372.001.

[115] M. F. Perutz, “Preparation of Haemoglobin crystals,” J. Cryst. Growth, vol. 2, no. 1, Feb. 1968, doi: 10.1016/0022-0248(68)90071-7.

[116] M. F. Perutz et al., “Three-dimensional Fourier Synthesis of Horse Oxyhaemoglobin at 2.8 Å Resolution : (I) X-ray Analysis,” Nature, vol. 219, no. 5149, pp. 29–32, Jul. 1968, doi: 10.1038/219029a0.

[117] M. F. Perutz, “X-ray Analysis of Hemoglobin,” Science (80-.)., vol. 140, no. 3569, pp. 863–869, May 1963, doi: 10.1126/science.140.3569.863.

[118] G. M. Artmann, “Microscopic photometric quantification of stiffness and relaxation time of red blood cells in a flow chamber,” Biorheology, 1995, doi: 10.1016/0006-355X(95)00032-5.

[119] R. M. Hochmuth, K. L. Buxbaum, and E. A. Evans, “Temperature dependence of the viscoelastic recovery of red cell membrane,” Biophys. J., vol. 29, no. 1, pp. 177–182, 1980, doi: 10.1016/S0006-3495(80)85124-1.

[120] W. Doster and S. Longeville, “Microscopic diffusion and hydrodynamic interactions of hemoglobin in red blood cells,” Biophys. J., vol. 93, no. 4, pp. 1360–1368, 2007, doi: 10.1529/biophysj.106.097956.

[121] Y. Kim, H. Shim, K. Kim, H. J. Park, S. Jang, and Y. K. Park, “Profiling individual human red blood cells using common-path diffraction optical tomography,” Sci. Rep., vol. 4, Oct. 2014, doi: 10.1038/srep06659.

[122] J. Wang, D. Bratko, and A. Luzar, “Probing surface tension additivity on chemically heterogeneous surfaces by a molecular approach,” vol. 108, no. 16, 2011, doi: 10.1073/pnas.1014970108/-/DCSupplemental.

[123] G. M. Artmann, “Habilitation: Methodische und experimentelle Beiträge zur Analyse der Ruheform, der Verformung und der Integrität humaner Erythrozyten.,” TU Ilmenau & Research Centre Juelich, Germany, Aachen, 1998.

[124] V. N. Uversky and A. V. Finkelstein, “Life in phases: Intra-and inter-molecular phase transitions in protein solutions,” Biomolecules, vol. 9, no. 12. MDPI AG, Dec. 01, 2019, doi: 10.3390/biom9120842.

[125] N. J. Greenfield, “Using circular dichroism spectra to estimate protein secondary structure,” Nat. Protoc., vol. 1, no. 6, pp. 2876–2890, Jan. 2007, doi: 10.1038/nprot.2006.202.

[126] V. Conti Nibali et al., “Wrapping Up Hydrophobic Hydration: Locality Matters,” J. Phys. Chem. Lett., vol. 11, no. 12, pp. 4809–4816, Jun. 2020, doi: 10.1021/acs.jpclett.0c00846.

[127] E. M. Adams et al., “Local Mutations Can Serve as a Game Changer for Global Protein Solvent Interaction,” JACS Au, vol. 1, no. 7, pp. 1076–1085, Jul. 2021, doi: 10.1021/jacsau.1c00155.

[128] S. G. Gevorkian, A. E. Allahverdyan, D. S. Gevorgyan, and C. K. Hu, “Glassy state of native collagen fibril?,” EPL, vol. 95, no. 2, Jul. 2011, doi: 10.1209/0295-5075/95/23001.

[129] S. G. Gevorkian, A. E. Allahverdyan, D. S. Gevorgyan, and C.-K. Hu, “Thermal-induced force release in oxyhemoglobin,” Sci. Rep., vol. 5, no. 1, Oct. 2015, doi: 10.1038/srep13064.

[130] Morozov V. and T. Morozova, “Viscoelastic Properties of Protein Crystals: Triclinic Crystals of Hen Egg White Lysozyme in Different Conditions,” Biopolymers, vol. 20, pp. 451–467, 1981.

[131] V. Kapil, C. Schran, A. Zen, J. Chen, C. J. Pickard, and A. Michaelides, “The first-principles phase diagram of monolayer nanoconfined water,” Nature, vol. 609, no. 7927, pp. 512–516, 2022, doi: 10.1038/s41586-022-05036-x.

[132] J. Ahlers et al., “The key role of solvent in condensation: Mapping water in liquid-liquid phase-separated FUS,” Biophys. J., vol. 120, no. 7, pp. 1266–1275, Apr. 2021, doi: 10.1016/j.bpj.2021.01.019.

[133] W. J. Ma, C. K. Hu, “Physical mechanism for biopolymers to aggregate and maintain in non-equilibrium states,” Sci. Rep., vol. 7, no. 1, Dec. 2017, doi: 10.1038/s41598-017-03136-7.

[134] S. Pezzotti, B. König, S. Ramos, G. Schwaab, and M. Havenith, “Liquid-Liquid Phase Separation? Ask the Water!,” J. Phys. Chem. Lett., vol. 14, no. 6, pp. 1556–1563, Feb. 2023, doi: 10.1021/acs.jpclett.2c02697.

[135] T. Bartfai, B. Conti, “Fever,” TheScientificWorldJournal, vol. 10. pp. 490–503, Mar. 16, 2010, doi: 10.1100/tsw.2010.50.

[136] C. M. Blatteis, “Endotoxic fever: New concepts of its regulation suggest new approaches to its management,” Pharmacol. Ther., vol. 111, no. 1, Jul. 2006, doi: 10.1016/j.pharmthera.2005.10.013.

[137] M. M. Diver, J. V. Lin King, D. Julius, and Y. Cheng, “Sensory TRP Channels in Three Dimensions,” Annu. Rev. Biochem., vol. 91, no. 1, pp. 629–649, Jun. 2022, doi: 10.1146/annurev-biochem-032620-105738.

[138] D. A. Yarmolinsky, Y. Peng, L. A. Pogorzala, M. Rutlin, M. A. Hoon, and C. S. Zuker, “Coding and Plasticity in the Mammalian Thermosensory System,” Neuron, vol. 92, no. 5, Dec. 2016, doi: 10.1016/j.neuron.2016.10.021.

[139] R. J. Schepers and M. Ringkamp, “Thermoreceptors and thermosensitive afferents,” Neuroscience and Biobehavioral Reviews, vol. 33, no. 3. pp. 205–212, Mar. 2009, doi: 10.1016/j.neubiorev.2008.07.009.

[140] M. Tominaga, M. J. Caterina, A. B. Malmberg, T. A. Rosen, H. Gilbert, and K. Skinner, “The Cloned Capsaicin Receptor Integrates Multiple Pain-Producing Stimuli the molecular entities at the nociceptor terminal that detect noxious signals and transduce this information into membrane depolarization events,” 1998.

[141] M. J. Caterina, A. Leffler, K. R. Petersen-Zeitz, M. M. Koltzenburg, A. I. Basbaum, and D. Julius, “Impaired Nociception and Pain Sensation in Mice Lacking the Capsaicin Receptor,” Science (80-.)., vol. 288, pp. 306–313, Apr. 2000.

[142] H. Xu et al., “TRPV3 is a calcium-permeable temperature-sensitive cation channel,” Nature, vol. 418, no. 6894, pp. 181–186, Jul. 2002, doi: 10.1038/nature00882.

[143] B. Nilius and T. Voets, “Channelling cold reception,” Nature, vol. 448, no. 7150, Jul. 2007, doi: 10.1038/448147a.

[144] J. Vriens, B. Nilius, and T. Voets, “Peripheral thermosensation in mammals,” Nat. Rev. Neurosci., vol. 15, no. 9, Sep. 2014, doi: 10.1038/nrn3784.

[145] D. D. McKemy, W. M. Neuhausser, and D. Julius, “Identification of a cold receptor reveals a general role for TRP channels in thermosensation,” 2002. [Online]. Available: www.nature.com.

[146] S. Brauchi, P. Orio, and R. Latorre, “Clues to understanding cold sensation: Thermodynamics and electrophysiological analysis of the cold receptor TRPM8,” 2004. [Online]. Available: www.pnas.orgcgidoi10.1073pnas.0406773101.

[147] R. Paricio-Montesinos et al., “The Sensory Coding of Warm Perception,” Neuron, vol. 106, no. 5, pp. 830–841.e3, Jun. 2020, doi: 10.1016/j.neuron.2020.02.035.

[148] T. Voets, G. Droogmans, U. Wissenbach, A. Janssens, V. Flockerzi, and B. Nilius, “The principle of temperature-dependent gating in cold-and heat-sensitive TRP channels,” 2004. [Online]. Available: www.nature.com/nature.

[149] R. Latorre, C. Zaelzer, and S. Brauchi, “Structure-functional intimacies of transient receptor potential channels,” Q. Rev. Biophys., vol. 42, no. 3, pp. 201–246, Aug. 2009, doi: 10.1017/S0033583509990072.

[150] J. Lei, R. U. Yoshimoto, T. Matsui, M. Amagai, M. A. Kido, and M. Tominaga, “Involvement of skin TRPV3 in temperature detection regulated by TMEM79 in mice,” Nat. Commun., vol. 14, no. 1, Dec. 2023, doi: 10.1038/s41467-023-39712-x.

[151] B. Liu, K. Hui, and F. Qin, “Thermodynamics of Heat Activation of Single Capsaicin Ion Channels VR1,” 2003.

[152] S. Chowdhury, B. W. Jarecki, and B. Chanda, “A molecular framework for temperature-dependent gating of ion channels,” Cell, vol. 158, no. 5, pp. 1148–1158, Aug. 2014, doi: 10.1016/j.cell.2014.07.026.

[153] E. A. Tansey, C. D. Johnson, “Recent advances in thermoregulation,” Adv. Physiol. Educ., vol. 39, no. 3, pp. 139–148, Sep. 2015, doi: 10.1152/advan.00126.2014.

[154] A. Clarke, P. Rothery, “Scaling of body temperature in mammals and birds,” Funct. Ecol., vol. 22, no. 1, pp. 58–67, Feb. 2008, doi: 10.1111/j.1365-2435.2007.01341.x.

[155] M. F. Perutz, A. J. Wilkinson, M. Paoli, and G. G. Dodson, “THE STEREOCHEMICAL MECHANISM OF THE COOPERATIVE EFFECTS IN HEMOGLOBIN REVISITED,” 1998. [Online]. Available: www.annualreviews.org.

[156] J. Baldwin, C. Chothia, “Haemoglobin: The structural changes related to ligand binding and its allosteric mechanism,” J. Mol. Biol., vol. 129, no. 2, pp. 175–220, Apr. 1979, doi: 10.1016/0022-2836(79)90277-8.

[157] D. C. Willford, E. P. Hill, “Modest effect of temperature on the porcine oxygen dissociation curve,” Respir. Physiol., vol. 64, no. 2, pp. 113–123, May 1986, doi: 10.1016/0034-5687(86)90035-6.

[158] R. E. Weber, K. L. Campbell, “Temperature dependence of haemoglobin–oxygen affinity in heterothermic vertebrates: mechanisms and biological significance,” Acta Physiol., vol. 202, no. 3, pp. 549–562, Jul. 2011, doi: 10.1111/j.1748-1716.2010.02204.x.

[159] R. M. Bowden, R. T. Paitz, “Temperature fluctuations and maternal estrogens as critical factors for understanding temperature-dependent sex determination in nature,” J. Exp. Zool. Part A Ecol. Integr. Physiol., vol. 329, no. 4–5, Apr. 2018, doi: 10.1002/jez.2183.

[160] E. L. Charnov, J. Bull, “When is sex environmentally determined?,” Nature, vol. 266, no. 5605, Apr. 1977, doi: 10.1038/266828a0.

[161] S. Hadži, J. Lah, “Origin of heat capacity increment in DNA folding: The hydration effect,” Biochim. Biophys. Acta - Gen. Subj., vol. 1865, no. 1, Jan. 2021, doi: 10.1016/j.bbagen.2020.129774.

[162] L. L. Xiong, M. A. Garrett, M. T. Buss, J. A. Kornfield, and M. G. Shapiro, “Tunable Temperature-Sensitive Transcriptional Activation Based on Lambda Repressor,” ACS Synth. Biol., vol. 11, no. 7, pp. 2518–2522, Jul. 2022, doi: 10.1021/acssynbio.2c00093.

[163] J. Boenigk, S. Jost, T. Stoeck, and T. Garstecki, “Differential thermal adaptation of clonal strains of a protist morphospecies originating from different climatic zones,” Environ. Microbiol., vol. 9, no. 3, pp. 593–602, Mar. 2007, doi: 10.1111/j.1462-2920.2006.01175.x.

[164] W. Yan, “An Equation for Modelling the Temperature Response of Plants using only the Cardinal Temperatures,” Ann. Bot., vol. 84, no. 5, Nov. 1999, doi: 10.1006/anbo.1999.0955.

[165] S. Bannai, Y. Sugita, Absorption spectra and reaction with haptoglobin of hemoglobin (α NO, β unliganded). Journal of Biological Chemistry 248(21),7527–7529(1973).

[166] Y. Yuan, T.J. Shen, P. Gupta, N.T. Ho, V. Simplaceanu, T. C. S. Tam, M. Hofreiter, A. Cooper, K.L. Campbell, & C. Ho, A Biochemical-Biophysical Study of Hemoglobins from Woolly Mammoth, Asian Elephant, and Humans. Biochemistry, 50(34), 7350(2011).

[167] R. N. Pittman, B. R. Duling, A new method for the measurement of percent oxyhemoglobin. Journal of Applied Physiology 38(2)(1975).

